# Convergent gain and loss of genomic islands drive lifestyle changes in plant-associated *Pseudomonas*

**DOI:** 10.1101/345488

**Authors:** Ryan A. Melnyk, Sarzana S. Hossain, Cara H. Haney

## Abstract

Host-associated bacteria can have both beneficial and detrimental effects on host health. While some of the molecular mechanisms that determine these outcomes are known, little is known about the evolutionary histories of pathogenic or mutualistic lifestyles. Using the model plant *Arabidopsis,* we found that closely related strains within the Pseudomonas fluorescens species complex promote plant growth and occasionally cause disease. To elucidate the genetic basis of the transition between commensalism and pathogenesis, we developed a computational pipeline and identified genomic islands that correlate with outcomes for plant health. One island containing genes for lipopeptide biosynthesis and quorum sensing is required for pathogenesis. Conservation of the quorum sensing machinery in this island allows pathogenic strains to eavesdrop on quorum signals in the environment and coordinate pathogenic behavior. We found that genomic loci associated with both pathogenic and commensal lifestyles were convergently gained and lost in multiple lineages through homologous recombination, possibly constituting an early step in the differentiation of pathogenic and commensal lifestyles. Collectively this work provides novel insights into the evolution of commensal and pathogenic lifestyles within a single clade of host-associated bacteria.

## Introduction

Host-adapted bacterial lifestyles range from mutualistic to commensal to pathogenic resulting in positive, neutral, or negative effects on host fitness, respectively^1^. Many of these intimate associations are the product of millions of years of co-evolution, resulting in a complex molecular dialogue between host and bacteria^2,3^. In contrast, horizontal gene transfer (HGT) can lead to rapid lifestyle transitions in host-associated bacteria through the gain and loss of virulence genes^4–6^. For example, the acquisition and loss of pathogenicity islands plays a key role in the emergence of enteropathogenic *E. coli* strains from commensal lineages and vice versa^5^. Similarly, a virulence plasmid transforms beneficial plant-associated *Rhodococcus* strains into pathogens, while strains without the plasmid revert to commensalism^4^. It is unclear if reversibility of lifestyles is common in other bacteria, or if acquisition of pathogenicity genes drives loss of genomic features associated with commensalism (or vice versa).

To further examine how changes in genome content might influence bacterial lifestyle, we focused on the *Pseudomonas fluorescens* (Pfl) species complex which contains both commensal strains and pathogens^7–13^. Pfl strains are enriched in close proximity to plant roots (the “rhizosphere”) relative to surrounding soil in diverse plants including the model plant *Arabidopsis thaliana*^9,14,15^. Single Pfl strains benefit *Arabidopsis* health by promoting lateral root formation, protecting against pathogens, and modulating plant immunity^9,16^. However, some Pfl strains cause diseases such as tomato pith necrosis^17^ and rice sheath rot^18^. Thus, we used the Pfl species complex in association with *Arabidopsis* to understand how strains shift along the symbiosis spectrum from pathogenic to commensal and how lifestyle might influence genome evolution.

Here we show that among several closely related *Pseudomonas* strains sharing >99% 16S rRNA identity, gain and loss of multiple genomic islands through homologous recombination can drive the transition from pathogenesis to commensalism. Using a novel high-throughput comparative genomics pipeline followed by reverse genetics, we found two unique sets of genomic features associated with predicted pathogenic and commensal strains. Evolutionary reconstruction indicates that gain and loss of these genomic features occurred multiple times, and that the gains and losses were mediated by homologous recombination of regions flanking conserved insertion sites. Collectively this work implicates interactions between homologous recombination and horizontal gene transfer as the primary drivers of lifestyle transitions in the rhizosphere.

## Materials and Methods

### Bacterial cultivation and quorum biosensor assay

All *Pseudomonas* wild-type and mutant strains in this study were routinely cultured in LB media at 28°C. Detailed information on all strains used in this study can be found in Table S1. The quorum sensing biosensor assay was carried out by restreaking single colonies of wild-type and mutant strains on LB with 1.5% agar next to a streak of *Chromobacterium violaceaum* CV026, then incubating as previously described^19,20^. For selected strains in the *P. brassicacearum* clade (as well as the Δ*luxI*_LPQ_ N2C3 mutant), we also performed a quantitative assessment of violacein production as previously described^21^. Briefly, each strain of interest was streaked three times next to a streak of *C. violaceum* CV026 on LB agar and incubated overnight. The next day, each CV026 streak was scraped off the plate using an inoculation loop and resuspended in 1 mL 10 mM MgSO_4_. 150 µL of the suspension was diluted into 10 mM MgSO_4_ prior to an absorbance reading at 764 nm to measure cell density. 764 nm was used to normalize cell density because violacein absorbs near the 600 nm wavelength normally used for optical density. The remaining suspension was centrifuged for 1 min at 6,000 rcf, the supernatant was removed, and the pellet was resuspended in 1 mL dimethyl sulfoxide and incubated for 30 min at room temperature. After incubation, samples were centrifuged at 15000 rcf for 15 mins. 500 µL of the supernatant was mixed with 500 µL 10 mM MgSO_4_ and the absorbance was measured at violacein’s absorbance peak of 585 nm. Relative violacein production was reported as the ratio of the OD_585_/OD_764_ measurements to account for differences in cell density.

### Gnotobiotic Arabidopsis root inoculation assays

*Arabidopsis thaliana* seeds of the Col-0 ecotype were sterilized with a 3-minute 70% ethanol treatment followed by 50% bleach for 10 minutes. Seeds were then washed thrice in sterile water and stored at 4°C in the dark for 48 hours prior to sowing on square plates with solid half-strength MS media containing no sucrose and 1% PhytoAgar. Seeds were sowed on the surface and the plates were sealed with MicroPore tape and stored vertically so that roots grew along the surface of the media. Seedlings were grown at 23°C under 100 µE cool white fluorescent lights and a 16 h light/8 h dark cycle. Roots were inoculated 5-7 days after sowing. Bacterial inocula were prepared by streaking out a freezer stock, then picking single colonies into an overnight 5 mL culture of LB. Aliquots from the overnight culture were centrifuged for 1 min at 6000xg, then resuspended in 10 mM MgSO_4_. The resuspensions were then diluted to an optical density measured at 600 nm of 0.1, followed by another 100-fold dilution into 10 mM MgSO_4_. 5 µL of this final dilution was used to inoculate along the primary root of each seedling. Plates were resealed with Micropore tape and returned to the growth chamber for 7-8 days, after which lateral roots were counted, primary roots were measured using scanned plates and ImageJ, and/or whole seedlings were weighed depending on the experiment. Boxplots for all quantitative measurements represent quartiles of the data with outliers discarded. Significance tests of mean values (with outliers included) were carried out using a two-sided unpaired T-test as implemented by the “stats.ttest_ind” function in SciPy (https://www.scipy.org). Lowercase letters are used on the boxplots to denote statistical significance below a threshold of p = 0.01.

### Gnotobiotic seed treatment assay for non-*Arabidopsis* plants

All seeds were sterilized with a 3-minute 70% ethanol treatment followed by 50% bleach for 10 minutes. 20-40 seeds were sown onto 80 mL of ½ strength MS media with 1% Phytoagar in cylindrical plant growth containers. 1 mL of overnight cultures of N2C3 and N2E3 in LB were centrifuged and washed three times in 10 mM MgSO_4_ before being diluted to an OD600 of 0.1. 1 mL of the bacterial suspension was dripped directly onto seeds immediately after sowing. Containers were returned to the growth chamber before being imaged at 8 or 15 days after inoculation.

### Gene deletions in *Pseudomonas* sp. N2C3

Deletion mutants were created using a double crossover methodology common for making deletions in Gram-negative bacteria^22^. Briefly, fragments of 800-900 bp flanking the gene of interest were amplified using PCR from genomic DNA. External primers had 5’ extensions adding a restriction site, while internal primers were designed to be in-frame with the gene of interest and had either another 5’ restriction site (Δ*luxI*) or a 21-bp linker (ΔSYP, ΔSYR, and Δ*luxR*). The fragments flanking the *luxI* gene were assembled using three-way ligation into the pNPTS138 vector (M. R. K. Alley, unpublished data) while the fragments flanking the remaining genes were assembled using overlap extension PCR into the pEXG2 suicide vector^23^. Primers for deletions can be found in Table S2.

Plasmids were transformed into chemically competent aliquots of the diaminopimelic acid (DAP) auxotroph *E. coli* WM3064 and plated on LB containing 0.3 mM DAP and the appropriate antibiotic (10 µg/mL gentamycin for pEXG2-derived vectors and 50 µg/mL kanamycin for pNPTS138-derived vectors). Individual colonies were picked into overnight cultures in LB with antibiotic and DAP. 1 mL of the transformed WM3064 overnight culture was washed 3x in LB to remove antibiotics before being centrifuged at 6,000xg with 1 mL of overnight culture of wild-type *Pseudomonas* sp. N2C3. The supernatant was decanted and the combined bacteria were resuspended in the liquid that remained (∼30 µL). This mixed bacterial suspension was spotted onto an LB plate and incubated at 28°C for 4-6 hours. After 4-6 hours, the spot was restreaked directly onto LB containing the appropriate antibiotic to isolate N2C3 transconjugants. We then restreaked strains again on the appropriate antibiotic before restreaking on no-salt LB with 10% sucrose for counterselection against the integrated plasmid.

We began our work using the pNPTS138 suicide vector which confers kanamycin resistance and sucrose sensitivity. However, we found that all *Pseudomonas* clones with an integrated pNPTS138 vector were still sucrose-resistant. Thus, for the Δ*luxI* mutant, we screened roughly ∼50 colonies to identify spontaneous mutants that lost the plasmid. This is consistent with a previous observation that the *sacB* gene was insufficiently expressed in *Pseudomonas aeruginosa* for lethality on sucrose-containing media, which led to the generation of pEXG2 which has increased expression of *sacB*^23^. Therefore, we used pEXG2 for all future deletions for stronger counterselection against *sacB*.

Once antibiotic-sensitive strains were recovered, colony PCR was performed using the upstream and downstream primers to distinguish wild-type revertants from deletion mutants. Deletion mutants were restreaked for purity, patched again on antibiotic-containing media to confirm plasmid loss, and verified again using colony PCR before being stored as a freezer stock. The ΔSYRΔSYP double mutant strain was constructed by introducing the ΔSYR deletion construct into the previously constructed ΔSYP mutant.

### 16S similarity

We extracted full-length 16S rRNA gene sequences for the strains in the *P. brassicacearum* clade. Using USEARCH to cluster 16S sequences at a threshold of 97% identity yielded a single cluster^24^. Increasing the USEARCH threshold to 99% yielded a second cluster containing two strains with more divergent 16S sequences (*Pseudomonas* sp. P97-38 and *Pseudomonas chlororaphis* GCA001023535), but the majority of sequences still mapped to a single cluster. This suggests that the diversity of this entire clade would be represented by a single OTU in amplicon-based sequencing studies.

### Computational Methods

A full description of the development, benchmarking and utilization of the PyParanoid pipeline, as well as details on comparative genomics and phylogenetic methods can be found in the Supplementary Information.

## Results

### A pathogen within a plant growth-promoting clade

To understand the emergence and phylogenetic distribution of plant-associated lifestyles, we focused on a well-characterized commensal strain with plant-beneficial activity and asked whether its closest cultured relatives also had beneficial effects on plant hosts. *Pseudomonas* sp. WCS365 robustly colonizes plant roots^25,26^, promotes growth^9^, and protects plants from soil-borne fungal pathogens^27^. Close relatives of WCS365 include an isolate from *Arabidopsis* (*Pseudomonas brassicacearum* NFM421)^28^, and isolates from a nitrate-reducing enrichment of groundwater (N2E2 and N2C3)^29,30^ (Figure 1A). Together, these 4 strains share nearly identical 16S rRNA sequences (>99.4% identity) and would be grouped into a single OTU in a community profile; however, it is unknown whether all members of this OTU share the beneficial antifungal and plant growth promotion abilities of WCS365.

**Figure 1.**
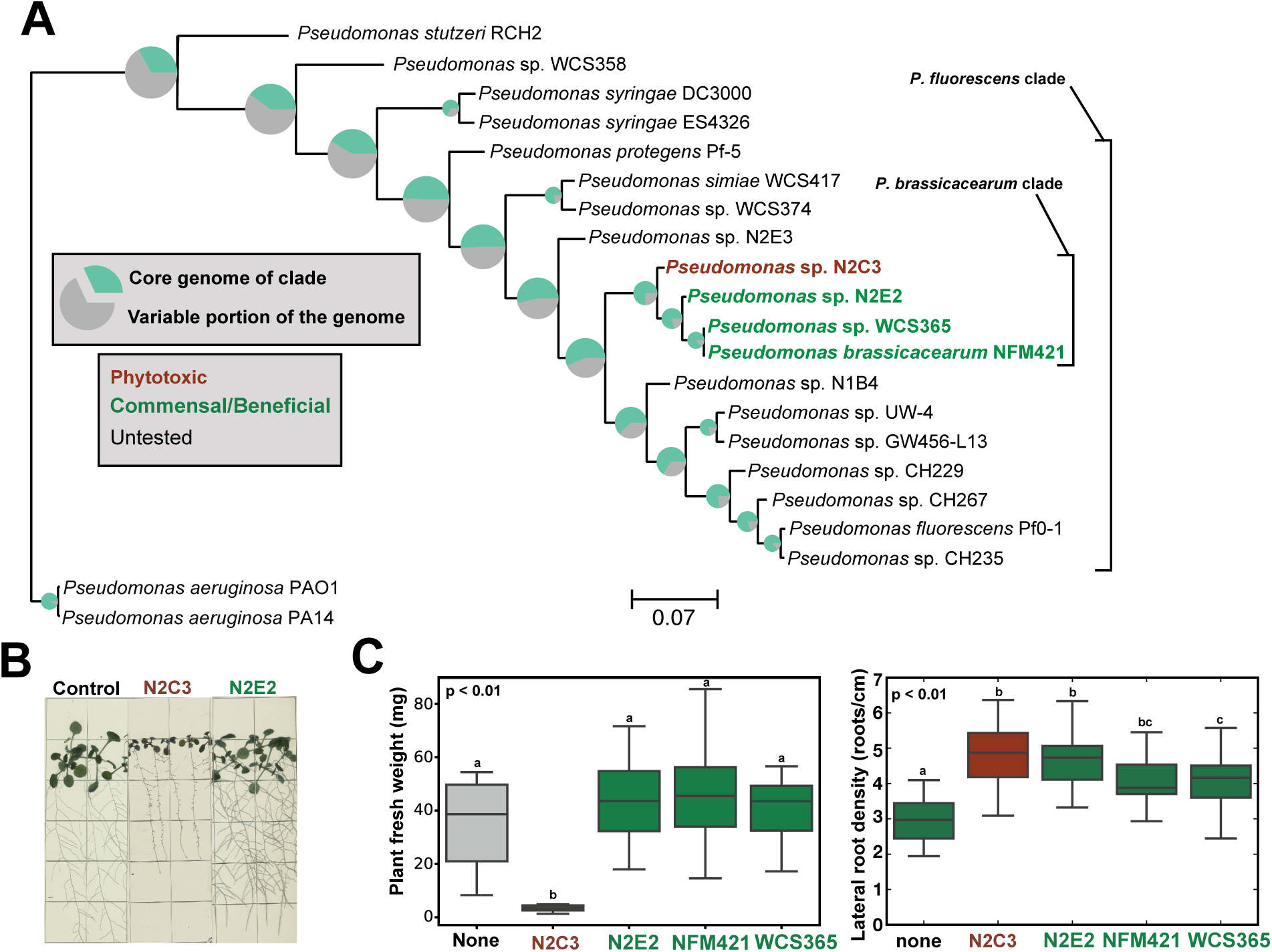
A pathogen within a clade of beneficial *Pseudomonas* spp. (**A**) A species tree of characterized *Pseudomonas* strains was generated using 217 conserved housekeeping genes. Strain names are color-coded to reflect plant-associated lifestyles based on prior reports in the literature. Pie charts at each node reflect the proportion of the genome that encodes “core” genes that are present in each member of the clade, showing that there can be large differences in gene content over short evolutionary distances. (**B**) Gnotobiotic *Arabidopsis* seedlings inoculated with N2C3, N2E2, or a buffer control. (**C**) Fresh weight of seedlings treated with 10 mM MgSO_4_, N2C3, N2E2, NFM421, and WCS365. Lateral root density of seedlings treated with 10 mM MgSO_4_, N2E2, NFM421, and WCS365. Lower-case letters indicate statistically significantly different (p < 0.01, 26 > n > 12) means as determined by a two-sided Student’s T-test for unpaired samples. Boxplots are quartile plots of the observed data. Data shown are from one of two independent experiments performed with similar results.

We tested whether these 4 closely related strains could promote plant growth. We found that in a gnotobiotic seedling assay where WCS365, NFM421 and N2E2 increased lateral root density and had no significant effect on fresh weight (Figure 1B-C), N2C3 caused significant stunting of primary root and rosette development (Figure 1B) and a significant reduction in fresh weight relative to mock-inoculated seedlings (Figure 1C). N2C3 also increased lateral root density, however, whether this is due to increased lateral root initiation or inhibition of primary root elongation is unclear (Figure 1B-C). Additionally, we found that N2C3 killed or stunted plants from the families *Brassicaceae* (kale, broccoli, and radish) and *Papaveroideae* (poppy), but had little to no effect on the *Solanaceae* (tomato and *Nicotiana benthamiana*) (Figure S1). Thus, unlike its close relatives that promote plant growth, N2C3 is a broad host range pathogen under laboratory conditions.

### A pathogenicity island found in plant-associated *P. fluorescens*

We reasoned that by comparing the genomic content of N2C3 to closely related commensal *P. fluorescens* strains and pathogenic *P. syringae* strains, we could identify the genetic mechanisms underlying pathogenicity or commensalism within this clade. The large number of sequenced genomes within the genus *Pseudomonas* made existing homolog detection methods (which scale exponentially) untenable for surveying the pangenome of the entire genus^31^. Therefore, we sought to develop a method that coupled fast but robust ortholog identification of a reference pangenome with a heuristic approach that generated binary homolog presence-absence data of the genes composing the reference pangenome for an arbitrarily large dataset.

In order to identify the genomic features associated with commensalism and pathogenesis, we built a bioinformatics pipeline called PyParanoid to generate the *Pseudomonas* reference pangenome on large genomic datasets. A detailed description of PyParanoid can be found in the Supplementary Information and an overview is shown in Figure S2. Briefly, PyParanoid uses conventional similarity clustering methods to identify the pangenome of a training dataset that includes phylogenetically diverse reference genomes and strains of experimental interest. The diversity of the training pangenome is then represented as a finite set of amino acid hidden Markov models (HMMs) which are then used in the second phase to catalog the pangenome content using computational resources that scale linearly (not exponentially) with the size of the dataset. The result of this pipeline is presence-absence data for a genome dataset that is not constrained by sampling density or phylogenetic diversity. This heuristic-driven approach enabled us to rapidly assign presence and absence of 24,066 discrete homology groups to 3,894 diverse genomes from the diverse *Pseudomonas* genus, assigning homology group membership to 94.2% of the 22.6 million protein sequences in our combined database (details in Supplemental Methods and Figure S2). The construction of the *Pseudomonas* pangenome database was accomplished using reasonable computational resources (roughly ∼230 core-hours on a single workstation). We also benchmarked PyParanoid on a series of test datasets against OrthoFinder2^32,33^. PyParanoid was much faster than OrthoFinder2 (2.7 hrs vs 36.3 hrs for a 120-strain dataset), but sacrificed no accuracy in homolog detection as determined by assessing the capture of a group of known single-copy genes, with both methods yielding a capture rate of 99.8% (details in Supplemental Methods and Figure S3).

Using the *Pseudomonas* reference pangenome, we searched for genes that were unique to the pathogenic N2C3 or its 3 closely related commensal relatives with plant-beneficial activity. We found that N2C3 contains a conspicuously large 143-kb island comprising 2.0% of the N2C3 genome that is not present in the other strains. The predicted functions of the genes are also consistent with a role in pathogenesis; the island features two adjacent large clusters of non-ribosomal peptide synthetase (NRPS) genes, as well as genes similar to the acyl-homoserine lactone (AHL) quorum sensing system prevalent in the Proteobacteria, which can play a role in virulence^34^ (Figure 2A). We designated this putative pathogenicity island the LPQ island (**l**ipo**p**eptide/**q**uorum-sensing). These clusters are very similar to genes involved in the production of cyclic lipopeptide pore-forming phytotoxins in *Pseudomonas syringae* spp. (syringopeptin and syringomycin), which have roles in virulence in many pathovars of *P. syringae*^35–37^. The genomic regions flanking the lipopeptide island are adjacent in the genome of N2E2 (Figure 2B) suggesting horizontal gene transfer (HGT) of the island. HGT has been reported for similar lipopeptide clusters in certain pathovars of *P. syringae*^38^.

**Figure 2.**
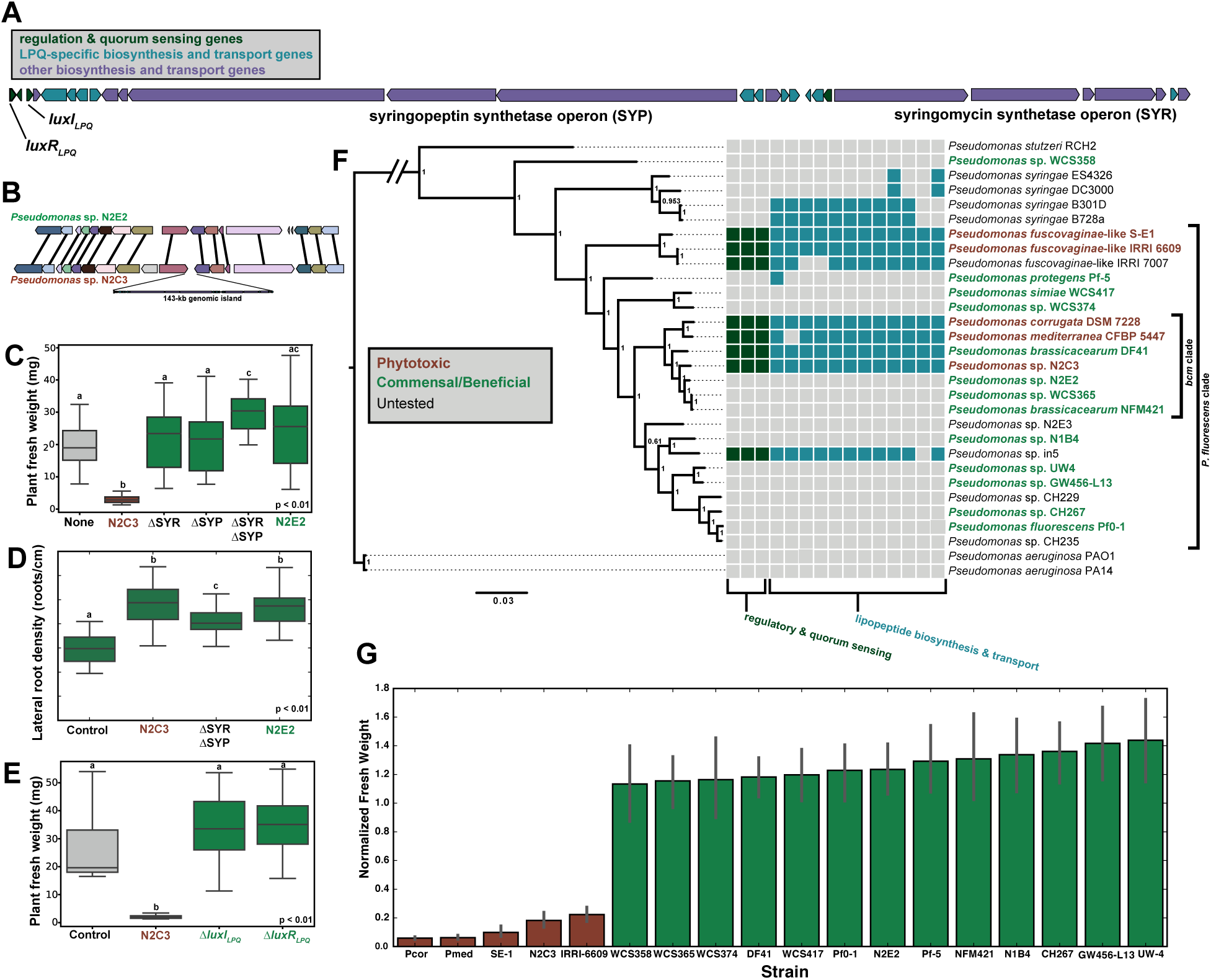
A genomic island necessary for pathogenesis. (**A**) A large genomic island encoding lipopeptide biosynthesis and quorum sensing genes (the LPQ island) is present in N2C3 but not in beneficial strains such as N2E2. (**B**) Diagram of the genomic island showing putative functions and locations of quorum sensing genes (*luxI*_*LPQ*_ – AHL synthase, *luxR*_*LPQ*_ – AHL-binding transcriptional regulator) and the lipopeptide biosynthesis clusters (syringomycin and syringopeptin). (**C**) Fresh weight of *Arabidopsis* treated with 10 mM MgSO_4_, wild-type N2C3, wild-type N2E2, N2C3 with a syringomycin cluster deletion (ΔSYR), N2C3 with a syringopeptin cluster deletion (ΔSYP) and a syringomycin and syringopeptin double deletion (ΔSYRΔSYP). Lower-case letters indicate statistically significant (p < 0.01, 20 > n > 15) differences as determined by a two-sided unpaired Student’s T-test. Boxplots are quartile plots of the observed data. Data shown are from one of two independent experiments performed with similar results. (**D**) Fresh weight of Arabidopsis treated with 10 mM MgSO_4_, wild-type N2C3, N2C3 with an AHL synthase deletion, (Δ*luxI*_*LPQ*_), and N2C3 with an AHL-binding transcriptional regulator deletion (Δ*luxR*_*LPQ*_). Lower-case letters indicate statistically significant (p < 0.01, 11 > n > 5) differences as determined by a two-sided unpaired Student’s T-test. Data shown are from one of two independent experiments performed with similar results. Boxplots are quartile plots of the observed data. (**E**) Lateral root density of seedlings treated with 10 mM MgSO_4_, wild-type N2C3, a syringomycin and syringopeptin double deletion (ΔSYRΔSYP), and wild-type N2E2. Lower-case letters indicate statistically significant (p < 0.01, 19 > n > 29) differences as determined by a two-sided unpaired Student’s T-test. Boxplots are quartile plots of the observed data. (**F**) Species tree showing the distribution of the island among divergent members of the *Pseudomonas* clade. Color coding of taxon labels summarizes the phenotypic data from Figure 2G. Colored squares indicate that an individual gene (as classified by PyParanoid) is present in a given taxon. Only 15 of the 28 genes in **B** (shown in green and teal in 2A, Data S1 for details) that are unique to the pathogenic strains are shown in **E**. The other 13 genes (shown in purple in **B**) are part of larger lipopeptide biosynthesis homolog groups that are not specific to the LPQ island. (**G**) Fresh weight of *Arabidopsis* seedlings treated with one of the 18 wild-type strains assayed in this study. Shown here are means of data from two independent experiments normalized to a 10 mM MgSO_4_ control (10 < n < 35). Error bars represent a 95% confidence interval for each mean using a non-parametric bootstrap of 1000 resamples.

In order to determine if the LPQ island is necessary for pathogenesis in the Pfl clade, we used reverse genetics to disrupt portions of the LPQ island in N2C3. We made clean deletion muants of gene clusters predicted to encode syringopeptin (ΔSYP - 73 kb) and syringomycin (ΔSYR - 39 kb), in addition to a mutant with both clusters deleted (ΔSYRΔSYP). We found that deletion of either cluster eliminated the N2C3 pathogenesis phenotype (Figure 2C). This is consistent with observations that both syringomycin and syringopeptin contribute to virulence in *P. syringae* B301D^35^. The ΔSYRΔSYP mutant elicited increased lateral root density compared to an untreated control (Figure 2D). We also generated knockouts of both the AHL synthase (LuxI_LPQ_) as well as the AHL-binding transcriptional regulator (LuxR_LPQ_). Both the Δ*luxI*_*LPQ*_ and Δ*luxR*_*LPQ*_ mutations abrogated the pathogenic phenotype (Figure 2E). These genetic results indicate that both lipopeptide biosynthesis and quorum sensing within the LPQ island are required for the pathogenicity of N2C3.

Because the LPQ island is necessary for pathogenesis in N2C3, we speculated that it may serve as a marker for pathogenic behavior in other *Pseudomonas* strains. We searched the PyParanoid database for other strains with genes contained within the 15 homology groups unique to the lipopeptide island (Table S3). While many of the lipopeptide biosynthesis-associated genes were found in a subset of *P. syringae* strains, the entire set of 15 genes including the quorum sensing system were found in three other species that contain bona fide plant pathogens (*P. corrugata, P. mediterranea* and *P. fuscovaginae* sensu lato) within the *P. fluorescens* clade as well as several strains with known antifungal activity (Figure 2F and Table S3). Genomic and genetic evidence from these three pathogenic species support a role for the LPQ island in pathogenesis in a variety of hosts, suggesting that the mechanism used by N2C3 to kill *Arabidopsis* may be conserved in divergent strains throughout the *P. fluorescens* clade^7,8,39–42^. Additionally, it was previously shown that the LPQ island is the source of antifungal cyclic lipopeptides in two other strains (DF41 and in5)^43,44^. Moreover, in *P. corrugata* and *P. mediterranea*, quorum sensing is found to have a role in modulating lipopeptide production, although it does not seem to affect lipopeptide production in DF41^41,42,45^. Collectively these data indicate that the LPQ island serves as a marker for plant pathogenic behavior and/or antifungal activity in diverse *Pseudomonas* spp.

To determine if the presence of the island predicted pathogenesis, we tested 14 additional isolates including five that contain the island and nine that do not. Of the five new isolates that contain the island, four (*P. mediterranea* CFBP 5447, *P. corrugata* DSM 7228, and *P. fuscovaginae*-like strains SE-1 and IRRI 6609) were originally isolated from diseased plant tissues and are capable of pathogenesis, whereas DF41 was isolated from canola root tips and is a commensal strain with antifungal activity (Table S1). The remaining nine new isolates that do not contain the island were a mix of plant-associated and environmental isolates. The four pathogenic isolates containing the island inhibited *Arabidopsis* to a similar degree as N2C3 (Figures 2F-G). On the other hand, DF41 did not inhibit *Arabidopsis* growth (Figures 2F-G) nor did any of the 9 new isolates that do not contain the island. The presence/absence of the LPQ island predicted pathogenic behavior in 17/18 (94%) of total wild-type isolates tested suggesting it may serve as a genetic marker for predicted pathogenic (LPQ+) or commensal (LPQ-) lifestyles within the *P. fluorescens* clade.

### Commensalism and pathogenesis are associated with multiple genomic features

Because deletion of the lipopeptide clusters converts N2C3 into a strain that increases lateral root density (ΔSYRΔSYP, Figure 2C-D), we considered whether presence or absence of the LPQ island alone might be sufficient to explain divergent bacterial lifestyles or if lifestyle changes are linked to additional genomic loci. To answer this question, we identified a broader monophyletic group of 85 strains with available genomes encompassing the *P. brassicacearum* clade, as well as the sister group containing the LPQ+ pathogens *P. corrugata* and *P. mediterranea* (hereafter the “*bcm* clade”, corresponds to the *P. corrugata* subgroup in ^31^). Together the *bcm* clade corresponds to the ‘*P. corrugata’* subgroup identified in other *Pseudomonas* phylogenomic studies and shares >97% 16S identity despite containing 8 different named species^31,46^. This group is broadly plant-associated, with 74 of the 85 genomes (87%) coming from strains isolated from plant tissue or rhizosphere (Table S1). Constraining our analysis to a phylogenetically narrow clade containing both pathogenic and commensal plant-associated bacteria allowed us to examine lifestyle transitions over a short evolutionary time.

To test if the pathogenicity island was correlated with the presence or absence of other elements of the variable genome, we performed a genome-wide association study (GWAS) in order to link the presence and absence of specific genes (based on PyParanoid data) with the predicted pathogenic phenotype (i.e. presence of the LPQ island). We utilized treeWAS, which is designed to account for the strong effect of population structure in bacterial datasets^47^. Using treeWAS, we identified 41 genes outside of the LPQ island which are significantly (p < 0.01) associated with the presence of the island based on three independent statistical tests (Table S4). 407 additional genes passed one or two significance tests, demonstrating that many genetic loci in the *bcm* clade are influenced by the presence of the LPQ island in the genome.

We explored the physical locations and annotations of the loci with significant associations with the LPQ island to identify clusters of genes with cohesive functional roles in plant-microbe interactions. Beyond the LPQ island we found 5 additional genomic loci: two are positively correlated with the LPQ island and 3 are negatively correlated with the LPQ island (Figure 3, Table S4). A subset of the genes significantly associated with the LPQ island are found in two small (<10kb) genetic clusters with unknown functions (putative pathogenicity islets I and II – PPI1 and PPI2) which are correlated with the presence of the LPQ island in validated pathogenic strains as well as LPQ+ genomes that have not been tested.

**Figure 3.**
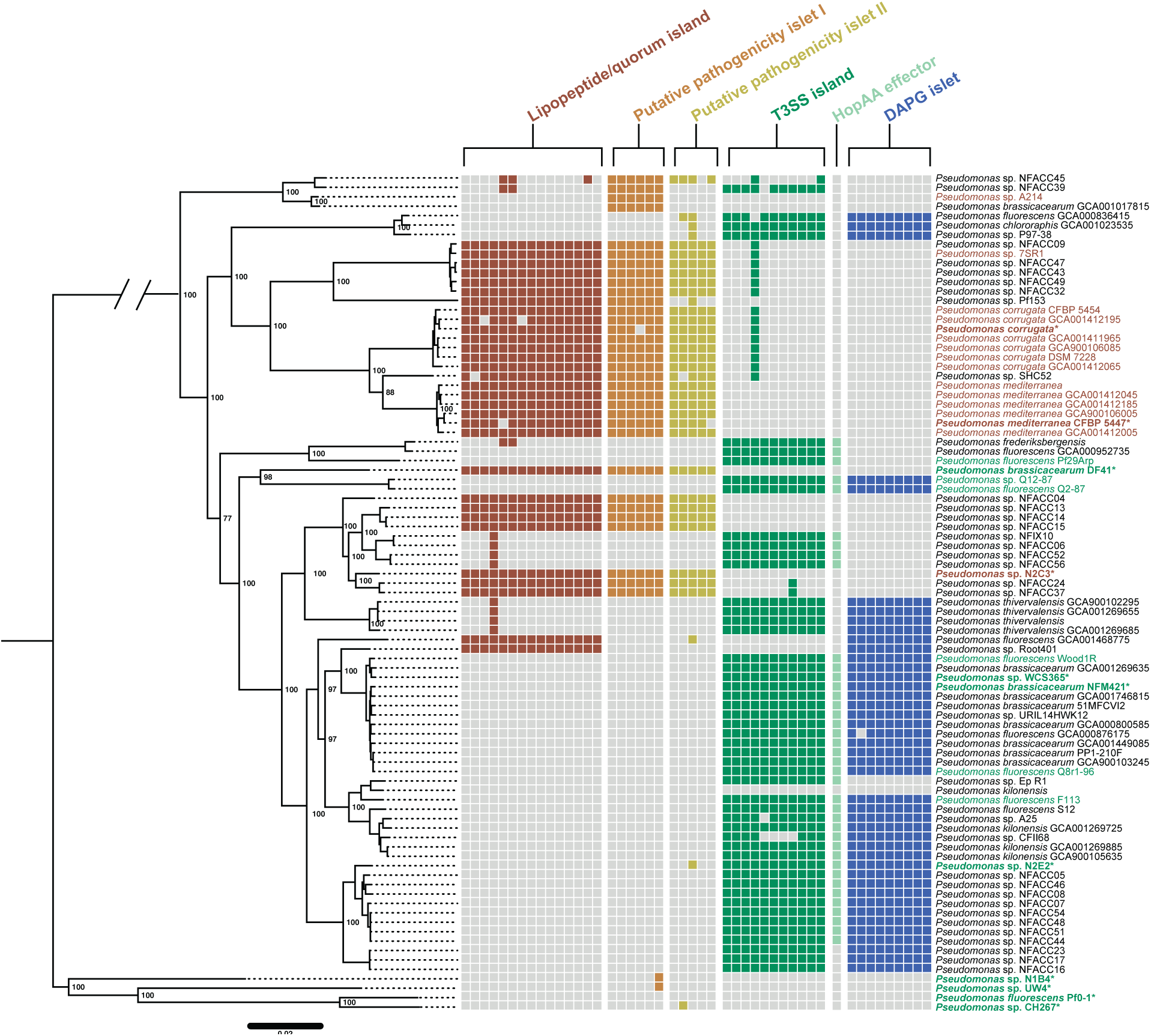
Polyphyletic distribution of pathogenic and commensal genomic islands within the *bcm* clade. A species tree was built for the *bcm* clade using a concatenated alignment of 2,030 conserved genes. Color-coded squares represent presence and absence of individual genes associated with each locus based on PyParanoid presence-absence data. In the case of both the LPQ and T3SS, only a subset of genes to the island are shown because both islands contain genes that are part of larger homolog groups not specific to either island. Bold taxon names highlight strains experimentally tested for commensal (green) or pathogenic (red) behavior (see Figure 2G). For strains that we did not test experimentally, we annotated rhizosphere lifestyle based on previous reports in the literature with the same color scheme. All six loci exhibit a highly polyphyletic distribution in extant strains suggesting that the pathogenic and commensal lifestyles have a complex evolutionary history in this clade.

The three loci that correlated with the absence of the LPQ island included a locus containing 28 genes encoding a type III secretion system (T3SS) and effectors (Table S4). This T3SS island is part of the broad Hrp family of T3SSs important for *P. syringae* virulence^2^. Despite their connotation as virulence genes, T3SSs can also be important for beneficial interactions like nodulation by rhizobia, providing a precedent for its occurrence in predicted commensal strains in the *bcm* clade^48^. The exact island identified here is found in commensal rhizosphere *bcm* clade strains Q8r1-96 and Pf29Arp where it is known as the Rop system and is necessary for the suppression of pathogen-and effector-triggered immunity (Figure 3)^49,50^. Moreover, the specific groups identified by PyParanoid as part of the Rop system are distinct from the Hrp genes of *P. syringae,* suggesting that T3SS genes in commensal rhizosphere bacteria have a distinct evolutionary history.

Many commensal strains in the *bcm* clade also have a single “orphaned” T3SS effector (T3SE) similar to the *P. syringae hopAA* gene (named *ropAA* in Q8r1-96)^49,51^. Commensal strains are also highly likely to contain a gene cluster for biosynthesis of diacetylphloroglucinol (DAPG), a well-studied and potent antifungal compound important for biocontrol of phytopathogens^52^. All 6 genetic loci (LPQ, PPI1, PPI2, T3SS, DAPG, and *hopAA,* Table S5) are polyphyletic, revealing a complex evolutionary history of lifestyle transitions within the *bcm* clade (Figure 3). Interestingly, these genomic features are largely restricted to the *bcm* clade, as a broader survey of the genus *Pseudomonas* and the *P. fluorescens* clade indicates only sporadic distribution of these genes (Figure S4). Collectively, this indicates that acquisition or loss of a pathogenicity island correlates with reciprocal gain and loss of genes that may be associated with commensalism within a clade of plant-associated of bacteria.

### Transitions between pathogenesis and commensalism arise from homologous recombination-driven genomic variation

To further understand the evolutionary history of the *bcm* clade, we searched for artifacts of the horizontal gene transfer (HGT) events that might cause the polyphyletic distribution of the 6 lifestyle-associated loci. For example, we might expect to see evidence of HGT such as genomic islands integrated at multiple distinct genomic locations or islands with a phylogenetic history very distinct from the core genome phylogeny. Additionally, we might find evidence of specific HGT mechanisms such as tRNA insertion sites, transposons, and plasmid-or prophage-associated genes^53,54^. We used the PyParanoid database to examine the flanking regions of each of the five islands and the *hopAA* gene. We detected each locus only in a single genomic context, with flanking regions conserved in all *bcm* genomes (Figures 4A and Figures S5-S10). These loci are not physically linked in any of the *bcm* genomes, suggesting that linkage disequilibrium of these loci is driven by ecological selection (“eco-LD”), not physical genetic linkage (Figure 4B and Figures S5-S10)^55^. Finally, there were no obvious genomic signatures of transposition, conjugation, transduction, or site-specific integration; all of which are commonly associated with horizontal gene transfer (HGT) of genomic islands^54,56^. Together, the absence of HGT signals and the conservation of the flanking regions signify homologous recombination of flanking regions as the primary mechanism driving gain or loss of the lifestyle-associated loci.

**Figure 4.**
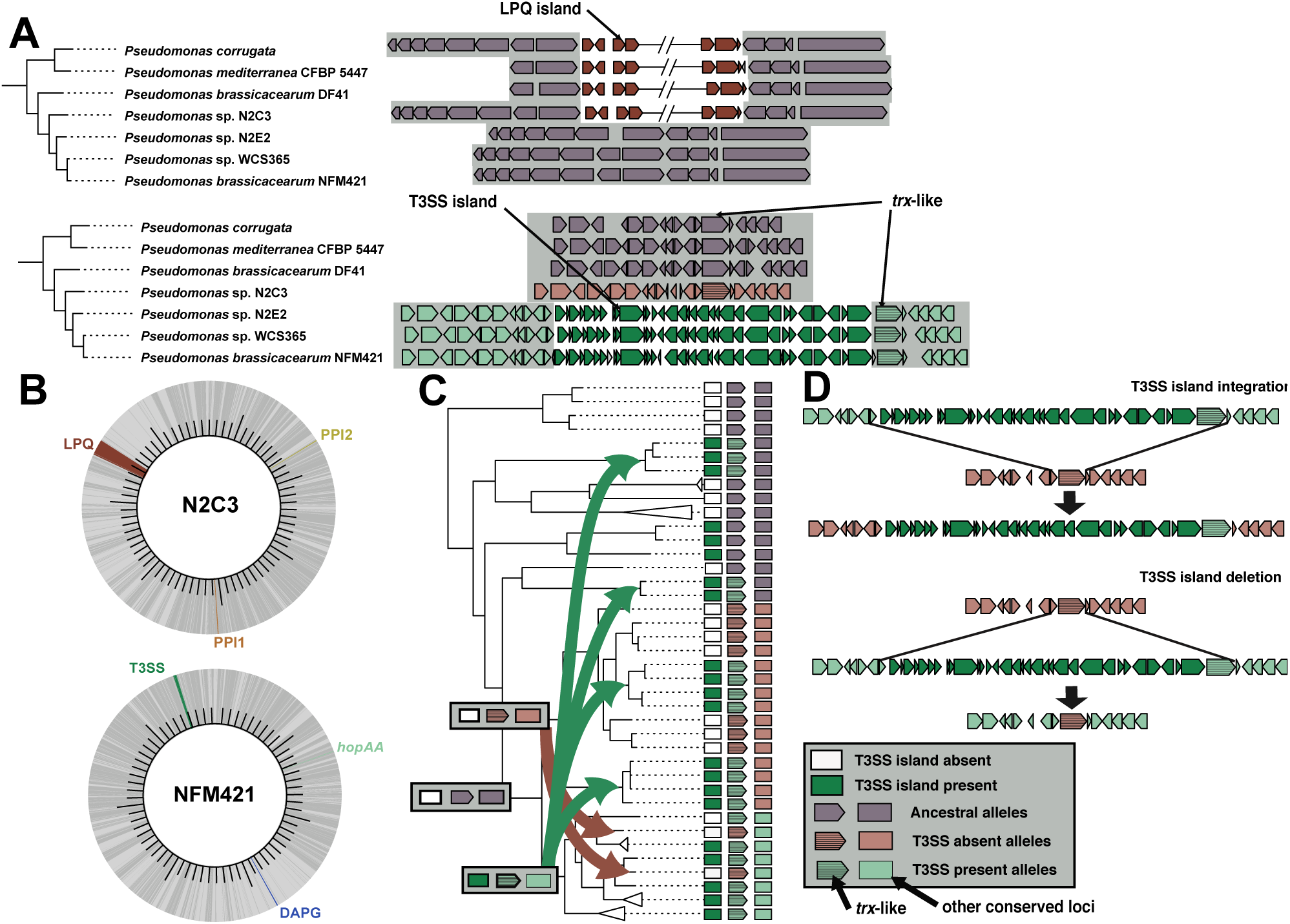
Phylogenomic evidence for independent island gain and loss through homologous recombination of flanking regions. (**A**) Insertion sites for each locus are conserved throughout the *bcm* species complex. The LPQ and T3SS island insertion sites are shown here for the 7 *bcm* clade strains that we characterized experimentally. All insertion sites for all 6 loci in the 85 strains can be found in Figures S5-S10. The tree shows the evolutionary relationship of the *bcm* strains. The grey background highlights that the conserved regions flanking both islands are adjacent in strains lacking the island. The location of the *trx-*like gene is shown adjacent to the T3SS which is used as a marker for recombination in Figure 4C. (**B**) Circular genome plots of two representative complete genomes from the pathogenic strain N2C3 and the beneficial strain NFM421. Darker grey regions of the genome represent the core genome of the *bcm* clade, while light grey denotes the variable genome. (**C**) A species phylogeny for the *bcm* clade adapted from Figure 3 to show evidence of homologous recombination of the T3SS island inferred through phylogenetic incongruencies between the species tree and the phylogeny of the *trx*-like gene (see Figure S13). Dark green rectangles denote taxa that have the T3SS island, while white rectangles represent taxa missing the island. Based on the phylogenetic evidence, we propose a parsimonious model for evolution of the T3SS island where gain of the T3SS island led to divergence of an ancestral lineage (purple arrow/rectangle) into a lineage with the T3SS present (light green arrow/rectangle) and a lineage with the T3SS absent (light red arrow/rectangle). However, subsequent gain (green arrows) or loss (red arrows) of the T3SS island has led to phylogenetic incongruence between the adjacent *trx-*like gene and the conserved loci used to construct the species tree (i.e. the *bcm* clade clonal phylogeny). (**D**) A diagram of the proposed homologous recombination mechanism leading to island gain (top) and loss (bottom). Both events lead to insertion of a *trx*-like allele with a distinct history from the surrounding conserved chromosomal loci.

Recombination events between distantly related strains can lead to incongruencies between gene and species phylogenies. To identify recombination events leading to island gain, we built phylogenies of the LPQ and T3SS islands and compared them to the species phylogeny. While the LPQ island phylogeny was largely congruent with the species phylogeny (Figure S11), the T3SS island had several incongruencies with the species tree (Figure S12). This indicates that recombination events leading to gain of the LPQ island were between closely related strains and are phylogenetically indistinguishable from clonal inheritance. In contrast, the history of the T3SS island shows evidence that the island was occasionally acquired from divergent donors.

Since the T3SS island’s history included several instances of recombination between distantly related donors and recipients, we reasoned that there might be signatures of such events in regions flanking the island. To test this hypothesis, we built phylogenies of conserved genes flanking the T3SS. For one gene downstream of the T3SS island integration site (annotated as ‘*trx-*like’, due to annotation as a thioredoxin-domain containing protein), we found that the gene tree was incongruent with the species tree, indicating horizontal gene transfer was prevalent in the history of the *trx-*like gene despite its conservation in all extant members of the *bcm* clade (Figure S13). By integrating the T3SS presence-absence data with the *trx*-like phylogeny and the species tree, we developed a model based on phylogenetic evidence that explains the origins of the T3SS island in extant *bcm* strains (Figures 4C, 4D and S13). Our model implicates homologous recombination between regions flanking genomic islands as the likely mechanism behind gain and loss of lifestyle-associated loci (Figure 4D). This provides an evolutionary mechanism underpinning the polyphyletic distribution of commensal-and pathogenic-associated islands and possibly strain lifestyle within closely related strains of *P. fluorescens*.

### Quorum interactions drive lipopeptide production and cooperative pathogenesis

Quorum sensing mechanisms are generally associated with monophyletic groups and their maintenance is thought to be enforced through kin selection^57^. However, the polyphyletic distribution of the LPQ island (Figure 4) suggests that distantly related LPQ+ strains might cooperate to induce pathogenesis to the exclusion of more closely-related LPQ-strains. If the *luxI*_*LPQ*_*/luxR*_*LPQ*_ system allows cooperation among distantly related LPQ+ strains, we would expect the system to be phylogenetically distinct from other AHL synthases and specifically associated with lipopeptide-producing strains within *Pseudomonas*. We found that LuxI_LPQ_ represented a monophyletic clade of *Pseudomonas* LuxI sequences as delineated using our *Pseudomonas* reference pangenome (Figure 5A). Furthermore, the presence of LuxI_LPQ_ had a positive correlation with all of the 14 other lipopeptide genes across the entire *Pseudomonas* clade (Figure 5B). While there are many lipopeptide-producing strains that lack LuxI_LPQ_ (mostly *P. syringae*), every strain that has LuxI_LPQ_ also has the entire LPQ island (Table S3). These *in silico* results conclude that LuxI_LPQ_ is specifically associated with cyclic lipopeptide-producing *Pseudomonas* spp. across the entire genus.

**Figure 5.**
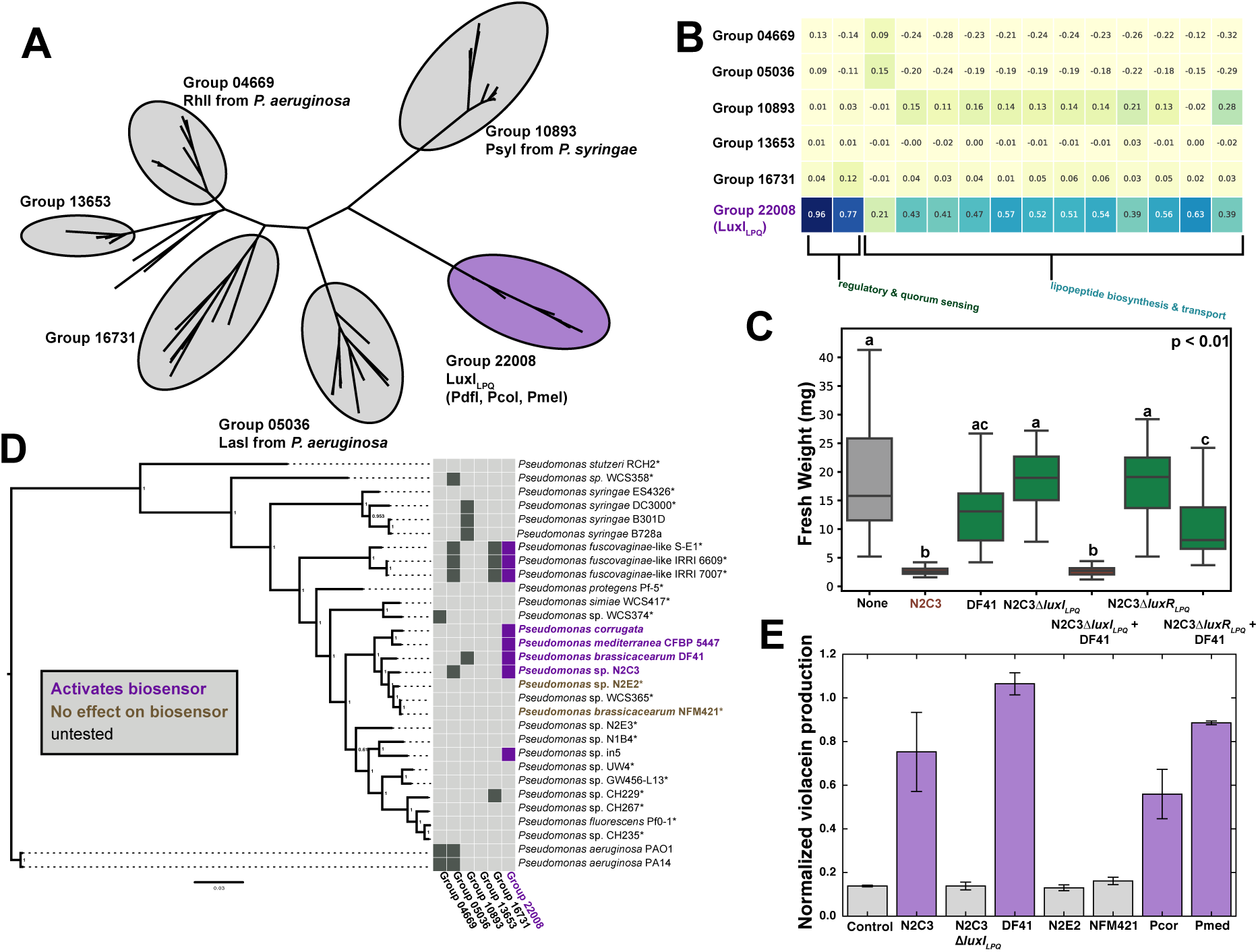
A lipopeptide-associated quorum sensing in divergent *Pseudomonas* spp. (**A**) Gene tree showing the phylogenetic relationship of the 6 AHL synthase homology groups identified by PyParanoid. Light gray squares indicate the absence of a gene and dark gray or purple squares show the presence of a gene. Group 22008 contains the LuxI_LPQ_ protein sequence from N2C3 as well as protein sequences from other LPQ+ strains (PdfI, PcoI, PmeI)^41,42,45^. (**B**) Correlation coefficients between each AHL synthase homology group and the 14 other LPQ island homology groups across the entire *Pseudomonas* clade showing that the presence of LuxI_LPQ_ is correlated with the capacity for lipopeptide biosynthesis across the entire *Pseudomonas* genus. LuxI_LPQ_ may serve as a common signal associated with lipopeptide production across diverse strains. (**C**) Fresh weight of seedlings inoculated with 10 mM MgSO_4_, wild-type N2C3, wild-type DF41, the AHL synthase deletion mutant N2C3Δ*luxI*_*LPQ*_ alone or co-inoculated with DF41, and the AHL-binding transcriptional regulator mutant N2C3Δ*luxR*_*LPQ*_ alone or co-inoculated with DF41. Lower-case letters indicate statistically significant (p < 0.01, 23 > n > 12) differences as determined by a two-sided unpaired Student’s T-test. Data shown are from one of two independent experiments performed with similar results. Boxplots are quartile plots of the observed data. (**D**) Species tree based on 217 housekeeping genes showing the distribution of 6 AHL synthase homology groups identified using PyParanoid. All strains with an asterisk were tested in a visual screen for induction of pigment production in the *C. violaceum* CV026 AHL biosensor strain, but only strains in the *bcm* clade with the LPQ island resulted in visible pigment production. Within the *bcm* clade, purple and brown taxon names show strains that were confirmed to induce or have no effect, respectively, on violacein production in a quantitative assay. (**E**) Barplot showing quantitative violacein production by *C. violaceum* CV026 by specified strains in the *bcm* clade. Data are the average of three biological replicates and error bars indicate the standard deviation (n = 3). Violacein production was normalized to total bacterial growth by calculating the ratio of OD_585_/OD_764_.

To test if the LuxI_LPQ_ homologs share the same signaling molecule, we co-inoculated *Arabidopsis* seedlings with DF41 (a non-pathogenic LPQ+ strain) and N2C3 Δ*luxI*_*LPQ*_ and Δ*luxR*_*LPQ*_ mutants, deficient in production of the AHL signal and signal perception, respectively. We found that DF41 restored pathogenicity of the non-pathogenic Δ*luxI*_*LPQ*_ AHL synthase mutant, indicating that it can provide an activating AHL signal *in trans*. However, DF41 did not restore pathogenicity of the Δ*luxR*_*LPQ*_ regulatory mutant (Figure 5C). Using a visual screen consisting of an AHL biosensor strain that produces the purple pigment violacein in response to short-chain AHL molecules, we found that all of the strains containing the LPQ island in the *P. brassicacearum* clade (*P. corrugata, P. mediterranea,* DF41 and N2C3) robustly elicited pigment production. The three *fuscovaginae-*like strains with the LPQ island (IRRI 6609, IRRI 7007, S-E1) did not activate the biosensor, nor did any of the strains without the island (0 out of 20) (Figure 5D)^20^. To confirm biosensor activity within the *bcm* clade, we utilized a quantitative assay for violacein production. We found that the four strains in the *bcm* clade containing the LPQ island (*P. corrugata, P. mediterranea,* DF41 and N2C3) increased violacein production, whereas two strains without the island (N2E2 and NFM421) and the N2C3 Δ*luxI*_*LPQ*_ strain failed to induce pigment production (Figure 5E). Reports from *P. corrugata, P. mediterranea, P. brassicacearum* DF41, and *P. fluorescens* in5 specifically implicate production of a C6-AHL molecule^41,42,45,58^, which is a strong inducer of the violacein-producing biosensor. Thus, the LPQ island has the capability to allow polyphyletic *Pseudomonas* spp. within the *bcm* clade to coordinate lipopeptide production and possibly pathogenesis through community C6-AHL levels with other LPQ+ strains.

## Discussion

Here we provide evidence that homologous recombination of a large pathogenicity island is associated with the transition between commensal and pathogenic lifestyles in a clade of plant-associated *Pseudomonas*. By focusing on the genetic variation within a single OTU (the *bcm* clade), we found that the presence of the pathogenicity island strongly predicted the presence or absence of 5 additional loci. Interestingly, the same island-based adaptations appear in multiple independent lineages, providing a compelling example of convergent gene gain and loss.

To understand the processes behind the convergent evolution within this clade, we gathered three independent lines of evidence that all support homologous recombination as the dominant mechanism of genomic island variation. The absence of signatures of horizontal gene transfer, the conservation of the flanking regions, and the incongruency of conserved flanking genes are all illustrative of homologous recombination. In particular, by reconciling the *trx*-like gene phylogeny with the species tree, we were able to identify specific instances where homologous recombination led to the gain or loss of the T3SS island (Figures 4, S13).

The genomic and phenotypic diversity of the *bcm* clade reveals the complexity inherent in studying the rhizosphere microbiome, particular when trying to link particular 16S sequences with functions in single strains. We found that labels like “commensal” and “pathogen” break down over short evolutionary distances within a well-studied clade of *Pseudomonas* spp. Moreover, we found that one strain (DF41) may function as a commensal or beneficial strain in isolation but might exacerbate the effects of bad actors through inter-strain quorum sensing. The role that the various loci play in the physiology and ecology of these strains (i.e. lifestyle) is still largely unclear. Thus, we emphasize that for the majority of strains in the *bcm* clade we are only predicting lifestyle; the actual effect on a plant host under different conditions must be determined empirically.

However, our core inference about lifestyle in the *bcm* clade based on the LPQ and T3SS islands relies substantially on experimentally validated evidence. The LPQ island is a known determinant of pathogenicity in the *bcm* clade^41,42^, whereas the T3SS has been found to be involved in suppressing host immunity in beneficial rhizosphere strains^49,50^. These two islands have a perfect anti-correlation in the *bcm* clade, with the T3SS occurring exclusively in beneficial or commensal strains, and the LPQ strains occurring only in strains described as pathogens with the exception of DF41 (Figure 2, Figure 3). DF41 had no negative effect on *Arabidopsis* in isolation (Figure 2), and has antifungal biocontrol activity^43^, thus appearing to be an authentic commensal. However, DF41 can make both the AHL signal molecule and lipopeptides, suggesting that regulation of the LPQ system in this strain is more complex than simple activation of lipopeptide biosynthesis by C6-AHL and may not be active in the rhizosphere.

Recent reports in DF41 (as well as the LPQ+ strain *P. fluorescens* in5 and *P. corrugata)* support a more complex regulatory mechanism by implicating two additional transcriptional regulators that have LuxR DNA-binding domains (but no AHL-binding domain)^45,58,59^. One of these genes, *rfiA*, is essential for lipopeptide production in DF41, while quorum sensing is not, demonstrating that the LPQ island in DF41 is regulated differently than in N2C3^45^. Identifying the signals and environmental conditions that activate lipopeptide production in different LPQ+ strains will be crucial for further elucidating the link between this island and antifungal and plant-pathogenic strain lifestyles.

The LPQ island has genome-wide implications for strains in the *bcm* clade, as we identified five loci in either positive or negative ecologically-driven linkage disequilibrium (“eco-LD”) with the LPQ island, implying epistasis and selection for one lifestyle or another. However, it is unclear how exactly microbe-mediated effects on the host might translate to microbial fitness in the rhizosphere and thus selection for the presence or absence of these loci. For example, do pathogenic strains outcompete commensal strains in a diseased plant? Furthermore, do recently diverging clades of pathogenic or commensal *bcm* strains even inhabit the same ecological niche? One possibility is that the LPQ island is a “niche-defining” evolutionary event that separates an incipient pathogen from its commensal predecessors, leading to further divergence^60^. Since the *bcm* clade contains pathogen to commensal transitions as well, the T3SS may have a similar niche-defining role, possibly manipulating immune responses of the host plant to favor other T3SS+ strains.

More broadly, our work provides evidence that epistatic genome-wide patterns in the pangenome have strong phenotypic implications for closely-related bacteria. These patterns may be difficult to spot when bacterial OTUs or populations are sparsely sampled. The recent increase of genomic data from many isolate populations makes GWAS and epistasis analyses a broadly powerful approach for identifying ecologically important loci that might not be identified using traditional genetics.

## Acknowledgements

We thank Dr. Clay Fuqua for providing the *Chromobacterium violaceum* CV026 biosensor strain, Dr. Teresa de Kievit and Dr. Ricardo Oliva for providing *Pseudomonas* isolates, and Dr. Adam Steinbrenner and Dr. Justin Meyer for critical reading of the manuscript. R.A.M. is a Simons Foundation Fellow of the Life Sciences Research Foundation. This work was also supported by an NSERC Discovery Grant (NSERC-RGPIN-2016-04121), Canada Foundation for Innovation, and Canada Research Chair grants awarded to C.H.H. The computational research was carried out with support provided by WestGrid and Compute Canada. R.A.M. and C.H.H. designed research and discussed results. R.A.M and S.S.H. performed experiments. R.A.M. wrote code and performed all bioinformatics analyses. R.A.M wrote the manuscript with input from C.H.H.

