## Supplemental Material for "Convergent gain and loss of genomic islands drive lifestyle changes in plant-associated *Pseudomonas*"

### Supplemental Methods

#### Overview of PyParanoid pipeline

A core task in any comparative genomics study is to identify homologous genes among a set of organisms. There are many methods available that address this task such as OrthoMCL[1], MultiParanoid[2], ProteinOrtho[3], and roary[4]. However, these methods do not effectively scale to genomic datasets of arbitrary size and phylogenetic divergence. Methods such as OrthoMCL that have "all-vs-all" sequence comparison steps have exponential increases in processing time[1,2]. Conversely, the roary algorithm eliminates highly redundant sequences to achieve faster run-times, but is constrained by use only on sets of closely related bacteria[4]. Modern microbial genomic databases are diverse collections of tens of thousands of genome sequences and thus must be pared down either in number or phylogenetic scope to identify homologs. For example, a recent phylogenomic study of the genus *Pseudomonas* identified upwards of 1200 publicly available genome sequences in the IMG database, but identified orthologs from a subset of only 166 genomes, presumably due to computational constraints[5]. We sought to develop methodology to allow us to incorporate all *Pseudomonas* genomes present in NCBI GenBank, which is even larger (>3800 strains annotated as *Pseudomonas* as of Fall 2017), into our comparative genomics study. The result of this effort is a new bioinformatic pipeline called PyParanoid.

PyParanoid is a set of scripts that knits together well-known bioinformatic tools into an efficient pipeline for homolog identification. PyParanoid consists of two phases: the first is an all-vs-all identification of the pangenome of a training dataset, while the second phase uses the pangenome to "annotate" presence or absence of each gene for each strain in the full dataset. The name PyParanoid reflects that the use of Python as the scripting language and the central role of the InParanoid pairwise homology detection algorithm in both phases[6].

During the first phase, protein FASTA files representing each of the strains in the training dataset are used as the input in an all-versus-all search using the computationally efficient sequence alignment algorithm DIAMOND[7], which is used in 'blastp' mode with a minimum score cut-off of 50. Initially, we hoped to use the raw bitscores from DIAMOND as input into the mcl clustering algorithm to detect groups of homologous protein sequences[8]. However, this resulted in many groups containing more than one sequence per genome indicating that paralogous families of genes were being merged into a single group. Therefore, we filter the pairwise similarity scores using the InParanoid algorithm, which use all-vs-all data to identify pairwise orthology relationships[6]. These pairwise relationships are then used as input to mcl, which leads to a large increase in the number of gene groups with a single member per strain, indicating robust detection of orthologous gene clusters. Highly similar (>95% identical) protein sequences are filtered out of each gene group using CD-HIT to mitigate bias due to overrepresented sets of strains[9]. The filtered groups are then aligned using MUSCLE using default parameters[10]. Each alignment is then used to build a profile hidden Markov model (HMM) using hmmbuild from the HMMER3 suite[11]. This set of HMMs represents the pangenome of the entire training dataset, minus "singletons" (i.e. a gene found in only a single strain).

We initially sought to use hmmscan/hmmsearch to exhaustively map the pangenome models (the "groups") to each additional strain in the test dataset using the HMM representation of each sequence profile. While this method scales linearly for each additional strain, the incremental computational cost is still very high. This is due to the fact that each HMM contains probabilities for all amino acids at each site. Therefore, we simplify the pangenome by generating consensus protein sequences for each model using hmmemit. These consensus sequences are combined into a single multi-FASTA file that was used for all-vs-all searches using DIAMOND against each additional strain. We again use InParanoid to filter the similarity scores into pairwise relationships between genes from the new strain and consensus sequence for the pangenome groups. These pairwise relationships are parsed to "annotate" each gene from each new strain with an existing homology group.

This "annotation" information along with the membership in the original gene clusters from the training dataset is used to construct the PyParanoid database, which consists of two tab-delimited text files, one containing presence/absence data and the other containing locus tags. Both files are indexed by strain and homology group ID, which is a five-digit number ("group_01234"). Additionally, the actual protein sequences for each homology group are extracted into multi-FASTA files for both the training and test datasets. Descriptions of each group are also generated by stripping annotation information from the headers of the FASTA input files, if available.

**Benchmarking PyParanoid**

We selected a set of 120 complete genomes from the 203 *Pseudomonas* genomes used to generate the 1st phase of the PyParanoid database to use for a benchmarking analysis. The 203 genomes were de-duplicated by generating a species tree and merging taxa separated by short branch lengths ($d < 0.005$). This yielded 132 strains, which we subsampled to 120 strains. We then randomized this list of strains to use as differently-sized datasets for benchmarking ($n = 5, 10, 20, 40, 60, 80, 120$). Each dataset contained the entire dataset from the smaller-sized datasets.

We selected Orthofinder2 for use as a benchmarking algorithm, due to its use of DIAMOND and mcl and its implementation in python[12,13]. To permit direct comparisons with PyParanoid, we used Orthofinder2 to only infer groups (i.e. using the '-og' option), not do phylogenetic paralog/ortholog delineation. Likewise, we ended our PyParanoid runs prior to aligning groups and building HMMs using the '–mode' option. For the first set of PyParanoid runs (PyP_core), we only used the 1st phase of PyParanoid which involves generating all-vs-all data for all strains in the dataset. For the second set (PyP_prop), we built models for the 40 strain PyParanoid dataset and propagated them to the larger benchmarking datasets ($n = 60, 80, 100, 120$). All analyses were carried out on the same compute cluster with each job using 8 cores and 32 Gb of memory.

We analyzed the computational resources required for these runs using the unix 'time' command. We calculated total CPU time as the sum of the 'user' and 'sys' values and total real time as the 'real' value returned by 'time'. For the PyP_prop runs, we summed the computational time from the PyP_core 40 strain run, the building of the models for PyP_core 40 strain, and the time required to propagate the models to the additional steps. We found that the PyP_core runs required drastically less total CPU time and total real time than OrthoFinder2 (Figure S3A-B). The PyP_prop runs required even less time than PyP_core, with computational resources barely increasing relative to dataset size over the samples tested. The slight increase in total real time for the PyP_prop run from $n = 40$ to $n = 60$ is due to the high input/output required for clustering, aligning, and HMM operations during the "build" phase of PyParanoid, which is normally only performed once per database.

To benchmark the capability of PyParanoid and OrthoFinder2 to capture known single-copy genes, we utilized the Gammaproteobacteria BUSCO (Benchmarking Universal Single-Copy Orthologs) database, which contains 452 single-copy genes[14,15]. We used the run_BUSCO script to annotate the 120 strains from the benchmarking dataset. Parsing the BUSCO results, we found that 410 of the 452 genes were found as a single copy in 115 or more of the 120 strains. We then extracted the locus tags for each of the 410 BUSCOs and used the OrthoFinder2, PyP_core, and PyP_build databases to assess which orthogroups corresponded to which BUSCO. For each BUSCO, we ranked the corresponding orthogroups by size and divided the largest orthogroup by the total number of tags in found in that BUSCO to determine the "capture rate". In other words, a capture rate of 1 indicates that 100% of the BUSCO tags were found in a single orthogroup. For each run, we averaged the capture rate across all 410 BUSCOs and reported the mean capture rates in Figure S3C. The capture rates were all very close to 1, indicating that across all three methods, one BUSCO corresponds to one orthogroup.

We also checked the reverse relationship, that is, if one orthogroup correspond to exactly one BUSCO. To do this, we calculated a specificity coefficient. Here, for each BUSCO, we took the top-ranked corresponding orthogroup and asked how many proteins in that group are there per strain. True orthologs with exactly one gene per strain have a value of 1. We averaged the BUSCO specificities for each run and found that all three methods were close to one (Figure S3D). Interestingly, as sample size increased, the specificity of orthogroups from OrthoFinder2 also increased, suggesting that duplicate genes are being added to the single-copy groups. Whether this reflects biological reality or is an artifact is unclear. Regardless, the BUSCO benchmarking indicates that PyParanoid performs similarly to OrthoFinder2 in identifying a set of conserved single-copy genes.

Finally, we calculated total orthologs (exactly one gene per strain) (Figure S3E), near orthologs (exactly one gene in >95% of strains) (Figure S3F), total groups (Figure S3G), and the fraction of the dataset clustered into orthogroups for each run (Figure S3H). The ortholog plots (Figure S3E-F) show that all 3 methods have similar trends in orthogroup detection. Interestingly, the number of total and near orthologs is larger for the PyParanoid methodologies than OrthoFinder in all samples tested. Again, whether this is reflective of biological reality or not is unclear. Figures S3G-H also demonstrate that OrthoFinder in general clusters datasets into fewer groups, but achieves higher coverage of datasets than the core PyParanoid methodology. Figure S3G-H also shows the key tradeoff of the PyParanoid propagation workflow. By restricting ortholog detection to the number of groups developed in the build phase (Figure S3G), the fraction of the dataset covered drops as sample size increases (Figure S3H). Overall, benchmarking confirmed that PyParanoid is an accurate orthogroup detection algorithm that sacrifices a modest amount of dataset coverage for a large increase in computational speed and scalability.

**Similar methods to PyParanoid.**

During the development of PyParanoid, two other high-throughput homolog detection pipelines were published that have some similarities to PyParanoid. panX utilizes DIAMOND and mcl to identify gene clusters, much like PyParanoid[16]. However it does not use any heuristics to reduce the size of the initial all-vs-all search or provide options for annotation using existing models[16]. Thus, using panX directly on large datasets requires substantial computational resources. panX also includes a graphical browser interface for analysis, which may complicate installation and use for novice users.

Like PyParanoid, DeNoGAP uses HMMs to represent individual gene families[17]. However, DeNoGAP is only available for UNIX-based systems. It also requires a functional SQLite installation and cannot be installed using a package manager. Unlike DeNoGAP and panX, PyParanoid is a lightweight python package and can be installed easily on any system from large compute clusters but also on local desktop and laptop machines running any operating system. The pipeline, as well as methods for analyzing and visualizing the database can be installed as a package using pip (pypi.python.org/pypi/PyParanoid). Secondly, PyParanoid is available at github.com/ryanmelnyk/PyParanoid. The github repository contains public troubleshooting and bug reporting threads, an installation guide and manual, as well as several iPython notebooks (in the "analysis_ipython" folder). The iPython notebooks illustrate how to use PyParanoid's built-in functions to carry out interactive analyses and visualization of a PyParanoid database.

While PyParanoid is exceptionally fast at calculating large pangenome databases, it lacks the sensitivity of conventional methods to detect rare homologs. As bacterial pangenomes are largely "open" and may contain an infinitely large pool of genes[18], the sequences unannotated by PyParanoid potentially represent a tail of thousands of low-abundance homology groups which may or may not be important for the phenotype of interest. Thus, PyParanoid is best for large-scale phylogenetic profiling, especially when looking at groups of genes, or large sets of strains. Selection of strains for the training set is also of utmost importance – if a strain of interest does not have a high-quality closed genome, it is recommended to be included in the training set since this will influence the gene models identified by PyParanoid. In this respect, DeNoGAP has a significant advantage over PyParanoid since it rigorously modifies, updates, and creates HMMs based on each incrementally added genome[17].

**Delineating the *Pseudomonas* clade**

Our initial goal in developing this pipeline was to build a cross-species phylogenomic database for thousands of strains within the genus *Pseudomonas*. However, we wanted to be confident that we were sampling a monophyletic group of bacteria, as taxonomic names given by submitters to public genome databases frequently do not reflect phylogenetic realities. Therefore, we downloaded 322 diverse and complete bacterial genomes from Ensembl Bacteria and used this as a training dataset for the first phase of PyParanoid to identify a set of gene families broadly conserved across all bacteria. These gene models could then be used in a manner similar to PhyloPhlan to rapidly generate phylogenies for various subsets of bacterial genomes[19]. We identified 5,235 gene families that were present in at least 10% of the training dataset and 122 "near-orthologs" that were present as a single copy in 95% of the training dataset.

We then used the 5,235 gene families from the 322 genomes in the training dataset to annotate 8,128 additional bacterial genome sequences (the "test dataset") using the second phase of PyParanoid. These genome sequences were sourced primarily from two public databases: all complete genomes from Ensembl Bacteria as well as all draft sequences designated as "*Pseudomonas*" from both Ensembl Bacteria and NCBI Genbank. We also added several *Pseudomonas* genomes from in-house sequencing projects which have since been uploaded to NCBI[20]. Of the 122 near-orthologs identified in the training dataset, 116.6 were found on average per genome in the test dataset, suggesting that our algorithm was correctly identifying orthologous genes conserved across Bacteria. We extracted the 122 gene families for all 8,450 genomes and generated a multiple sequence alignment by using hmmalign (http://hmmer.org/) to align each protein sequence to the HMM profile for that family. By aligning each protein sequence to a single HMM profile, the computational complexity of this step also scales linearly with the number of sequences unlike conventional multiple sequence alignment algorithms, which scale exponentially. After concatenating these alignments, we subsampled 30,000 random amino acid residues to generate an "all-bacteria" reference phylogeny using FastTree 2 with the JTT model of amino acid evolution[21]. Within the all-bacteria phylogeny we identified a node separating the Proteobacteria from the rest of the dataset. Returning to the PyParanoid database, we identified 217 near-orthologs conserved in the Proteobacteria which would allow for finer resolution when examining the *Pseudomonas* clade.

**Using PyParanoid to build the *Pseudomonas* pangenome database**

Within the all-bacteria phylogeny, 3,890 of the 3,891 genomes annotated as *Pseudomonas* resided in a single well-supported clade. This clade also included 4 strains annotated as *Azotobacter* which are located within the large *Pseudomonas* genus[5]. To improve resolution of the phylogenetic relationships within this clade, we extracted the 217 Proteobacteria-specific orthologs from the 3,894 *Pseudomonas* and *Azotobacter* genomes, finding 211.6 on average per strain. 5,000 random amino acid residues were subsampled without replacement (i.e. a single jackknife) from the aligned orthologs and used to generate a *Pseudomonas* phylogeny using FastTree 2. Subsets of strains from this alignment were used to construct some of the "species trees" depicted in this paper (Figures 1A, 2F, 4A, and 5D) with the exception of the tree in Figure 3 (see below). This tree was rooted to a clade of *Pseudomonas* that formed a well-supported, early-diverging branch in the all-bacteria phylogeny. Using this tree as a reference, we selected 203 completed genomes distributed broadly across the training dataset. Notably, we ensured that *Pseudomonas* sp. N2C3, *Pseudomonas* sp. N2E2, and *Pseudomonas brassicacearum* NFM421 (3 of the 4 strains of interest with finished genomes, Figure 1) were included in the training dataset.

The first phase of the PyParanoid pipeline identified 24,066 gene families which classified 98.0% of the 1.1 million sequences in the dataset. 293 gene families were found as a single copy in all strains ("true orthologs") and 1,784 "near orthologs" were found as a single copy in >95% of the 203 strains. For comparison, a recent study identified 794 orthologs conserved in *Pseudomonas* using a dataset of similar phylogenetic breadth but fewer strains (166 total genomes)[5].

In the second phase of PyParanoid, the 24,066 gene families were used to annotate the 3,691 remaining draft *Pseudomonas* genomes. These 24,066 families were sufficient to cover 93.2% of the protein sequences in the test dataset (20.0 of 21.5 million). Together the first and second phases of PyParanoid generated a *Pseudomonas* reference pangenome database which covered 94.2% of the 22.6 million protein sequences in the combined dataset. Using the BUSCO benchmarking method described earlier, we found that the entire dataset had a single-copy ortholog capture rate of 0.946 and an ortholog specificity of 1.001, showing that PyParanoid retains accuracy even with thousands of genomes, many of which are highly fragmented. This *Pseudomonas* pangenome database formed the entire basis of all comparative genomics analyses in this paper.

**Phylogeny of the *bcm* species complex**

We extracted 85 strains that corresponded to a well-supported monophyletic clade within the *Pseudomonas* tree containing species designated as *P. brassicacearum, P. corrugata, P. mediterranea* and others (hereafter called the *bcm* clade for simplicity). This clade corresponds to the '*P. corrugata*' subgroup in previous phylogenomic studies of the *P. fluorescens* clade and the entire *Pseudomonas* genus[5,22]. Using 4 other *Pseudomonas* spp. genomes as an outgroup, we generated a high-resolution alignment of 2,030 single copy orthologs shared by all 89 strains. This alignment consisted of 631,030 amino acid residues with 101,757 informative positions. By using ProtTest 3 on several 10,000 amino acid jackknifed samples from this alignment, we determined that the JTT model with empirical amino acid frequencies was the best model of amino acid evolution for this dataset according to the Akaike Information Criterion[23]. We then used RAxMLv8.2.9 to generate 20 maximum-likelihood trees and 100 independent non-parametric bootstraps using the JTT model with empirical frequencies under the gamma rate distribution[24]. The 100 bootstraps were drawn onto the topology of the best-scoring maximum-likelihood tree in order to generate the tree with node support values shown in Figure 3.

**Phylogeny for taxa tree**

To generate the tree for the entire *Pseudomonas* genus in Figure S4A, we made use of the alignment of 122 single-copy genes we generated for the bacteria-wide phylogeny described above. For computational feasibility, the alignment was randomly subsampled to 10,000 amino acid positions, ignoring sites that were highly gapped (>20%). FastTree v2.1.9 was used to build the phylogeny using default parameters. The phylogeny was rooted to a clade of *Pseudomonas* identified as an outgroup to all other *Pseudomonas* spp. in our previous bacteria-wide analysis. To more easily visualize this tree, we collapsed monophyletic clades with strong support (as determined by FastTree's local Shimodaira-Hasegawa test) that correspond with major taxonomic divisions identified by a recent phylogenomic survey of *Pseudomonas* spp[25]. This survey also used a similar outgroup choice and had a very similar topology to our tree.

To build the tree for the *Pseudomonas fluorescens* (*Pfl*) subclade seen in Figure S4B, we identified 1,873 orthologs specific to the *Pfl* clade found in >99% of all strains in the clade and then aligned them all to the HMMs generated by PyParanoid using hmmalign, prior to concatenation. This alignment had 581,023 amino acid positions, which we trimmed to 575,629 positions after masking sites with >10% of taxa with gaps. From this alignment, we randomly subsampled 120,000 sites for our final phylogenomic dataset. Using RAxMLv8.2.9, we inferred 20 independent trees under the JTT substitution model using empirical amino acid frequencies and selected the one with the highest likelihood. Support values were calculated through 100 independent bootstrap replicates under the same parameters. The tree was rooted to the *P. fragi* clade, which we identified as an outgroup from the *Pseudomonas*-wide phylogeny. Again, clades were collapsed and name by comparing our phylogeny to the Hesse *et al.*, 2018 tree, which also used *P. fragi* as an outgroup and had a similar topology to our tree[25].

**Identifying lifestyle-associated loci using treeWAS**

In order to identify other possible events of gene gain or loss linked with the lipopeptide island, we utilized the program treeWAS, which was recently developed to address the challenges specific to microbial genome-wide association studies (GWAS)[26]. Namely, by using phylogenetic information, treeWAS corrects for the effect of clonal microbial population structure. Therefore, we used the *bcm* phylogeny and a gene presence-absence matrix for the 85 *bcm* strains as the input to treeWAS. The phenotype information for treeWAS was encoded based on the presence of the LPQ island: putatively pathogenic if present, and putatively commensal if absent. Essentially, we were using treeWAS to look for genes associated with the LPQ island across the *bcm* clade. Using treeWAS, we generated significance values for associations using the simultaneous, subsequent and terminal tests (Data S2). We also generated coefficients of correlation between the presence of each individual orthology group and the lipopeptide island to determine whether the presence or absence of a given locus is associated with the LPQ island in conjunction with the significance scores reported from treeWAS. 60 of the 62 genes that were found to be significant in all three tests were found to be clustered in four genetic loci. 21 of these 60 genes were in the LPQ island, which is expected since this was how we defined the pathogenic phenotype. Eleven genes were found in two putative pathogenicity islets (PPI1 and PPI2) associated with presence of the island. 28 of the 60 genes were found in a type III secretion system (T3SS) island which was associated with the absence of the island. Exploring the annotations for the 407 other genes which passed one or two of the treeWAS significance tests revealed two other potentially beneficial loci: the diacetylphloroglucinol biosynthetic cluster (DAPG) and the orphaned *hopAA* effector. Locus tags from the *bcm* clade for all 6 loci identified using treeWAS can be found in Data S3.

**Concatenated alignment phylogeny of gene trees and tanglegrams**

For the LPQ and T3SS islands, homologs specific to each island were extracted across the entire dataset (i.e. all *Pseudomonas*), aligned using hmmalign, and concatenated into a single alignment, prior to phylogeny construction with FastTree 2. In both cases, the *bcm* clade sequences formed a monophyletic group. Thus, we removed taxa from each alignment to include only the *bcm* strains and an outgroup of a few *Pseudomonas* strains outside of the *bcm* clade. We then used ProtTest 3 to select the best protein evolution model for the lipopeptide and T3SS alignments which was JTT with empirical residue frequencies for both islands. RAxML was again used to build trees for these two alignments, drawing 500 independent non-parametric bootstraps onto the best tree from 50 independent maximum-likelihood trees. For the *trx*-like phylogeny, we used a nucleotide alignment and included two non-*bcm* clade *P. fluorescens trx*-like alleles as the outgroup. This alignment was used to generate a maximum-likelihood tree using FastTree 2 using a Jukes-Cantor model of nucleotide evolution.

To determine the degree of incongruence for the LPQ and T3SS islands relative to the species phylogeny, we generated pairs of rooted trees for both the locus of interest from *bcm* clade strains ("locus tree") and a 10,000-site jackknifed alignment from the *bcm* clade containing only the taxa where the locus of interest was present ("species tree"). Both trees were generated using the JTT model of amino acid evolution with empirical frequencies and by choosing the best-scoring tree from 20 independent maximum-likelihood trees. Support values were generated from 100 independent non-parametric bootstraps. The location of the root in the locus tree was determined using the tree with the non-*bcm* taxa from the previous paragraph, whereas the root of the species tree was determined using the *bcm* phylogeny in Figure 4. Rooted tree pairs were viewed and displayed in "tanglegram" format using Dendroscope 3[27].

For the AHL synthase multi-family tree (Fig. 5A) we used the AHL synthase Pfam (PF00765) to search the *Pseudomonas* reference pangenome. Six groups were identified and the individual protein sequences from all were aligned to the AHL synthase Pfam HMM using hmmalign. This alignment was used to construct a tree with FastTree2 which uses the JTT model by default for amino acid data.

**Data and materials availability:** All data is available in the manuscript or the Supplementary Materials. Source code for PyParanoid is available at https://github.com/ryanmelnyk/PyParanoid.

**Supplemental Figures**

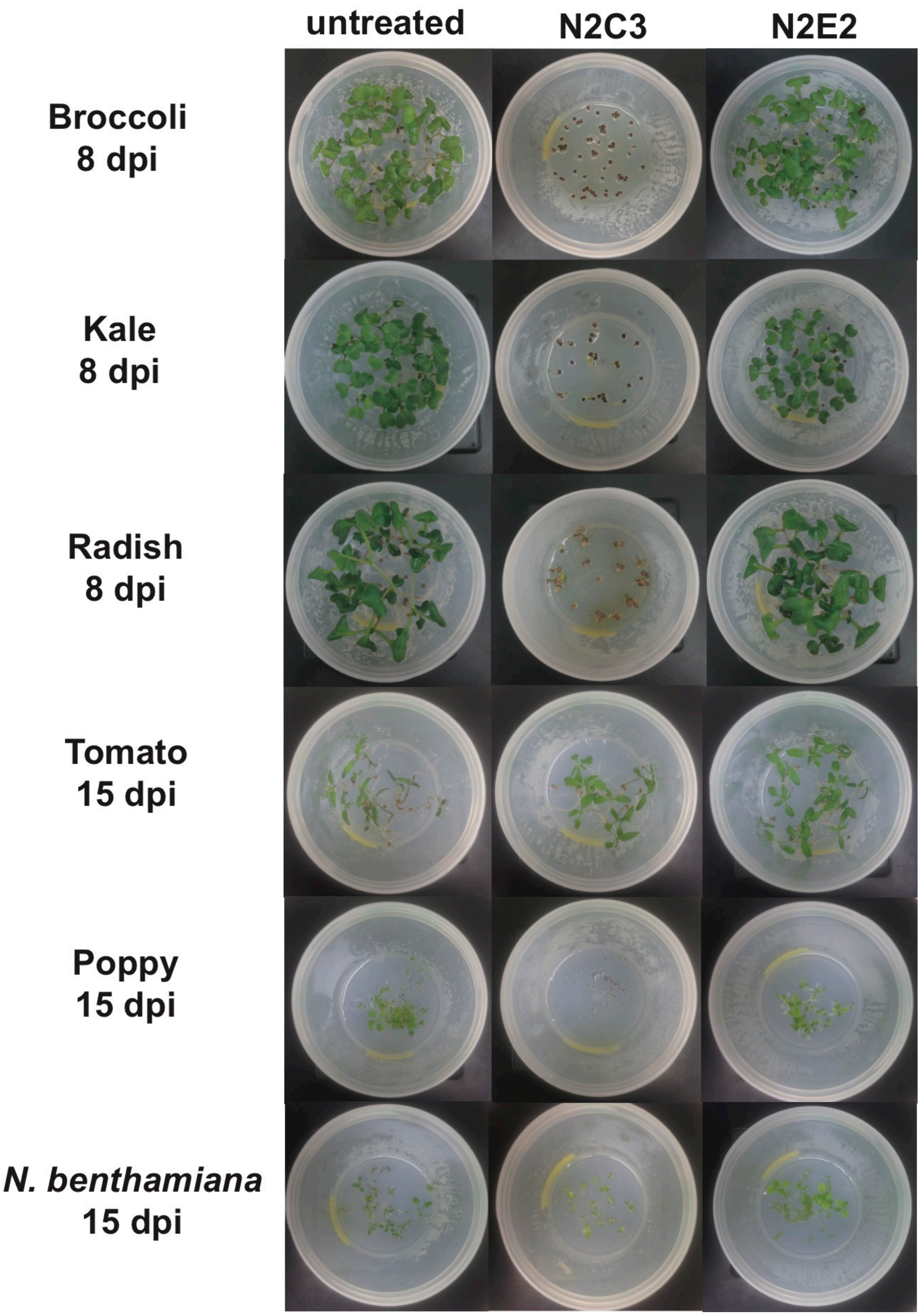


***Figure S1.*** N2C3 is capable of causing disease on diverse host plants under gnotobiotic conditions. Seeds from diverse plants were treated with a suspension of either N2C3 or N2E2. N2C3 inhibited germination of all seeds tested from the *Brassicaceae* (broccoli, kale, and radish) as well as poppy (*Papaveroidae*). N2C3 had no effect on tomato, but may cause slightly impaired growth of *Nicotiana benthamiana*. N2E2 treatment did not stunt plant growth under these conditions.

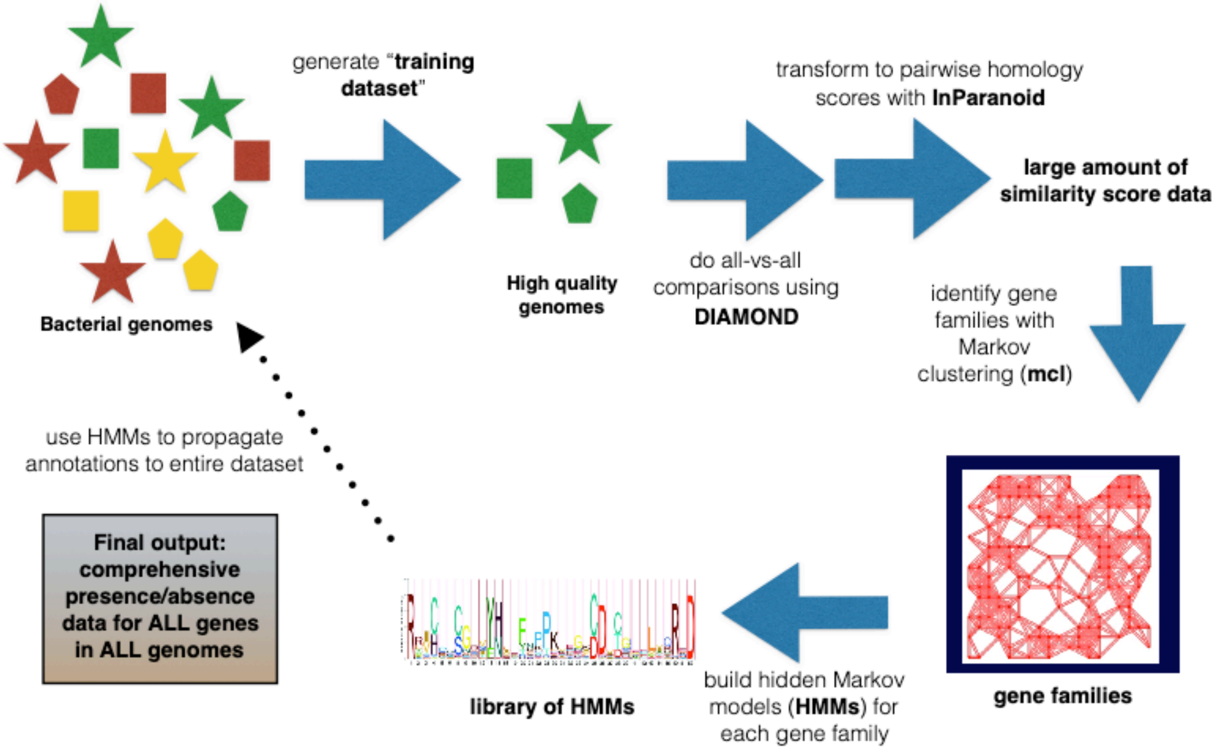


***Figure S2.*** Visual overview of the PyParanoid pipeline.

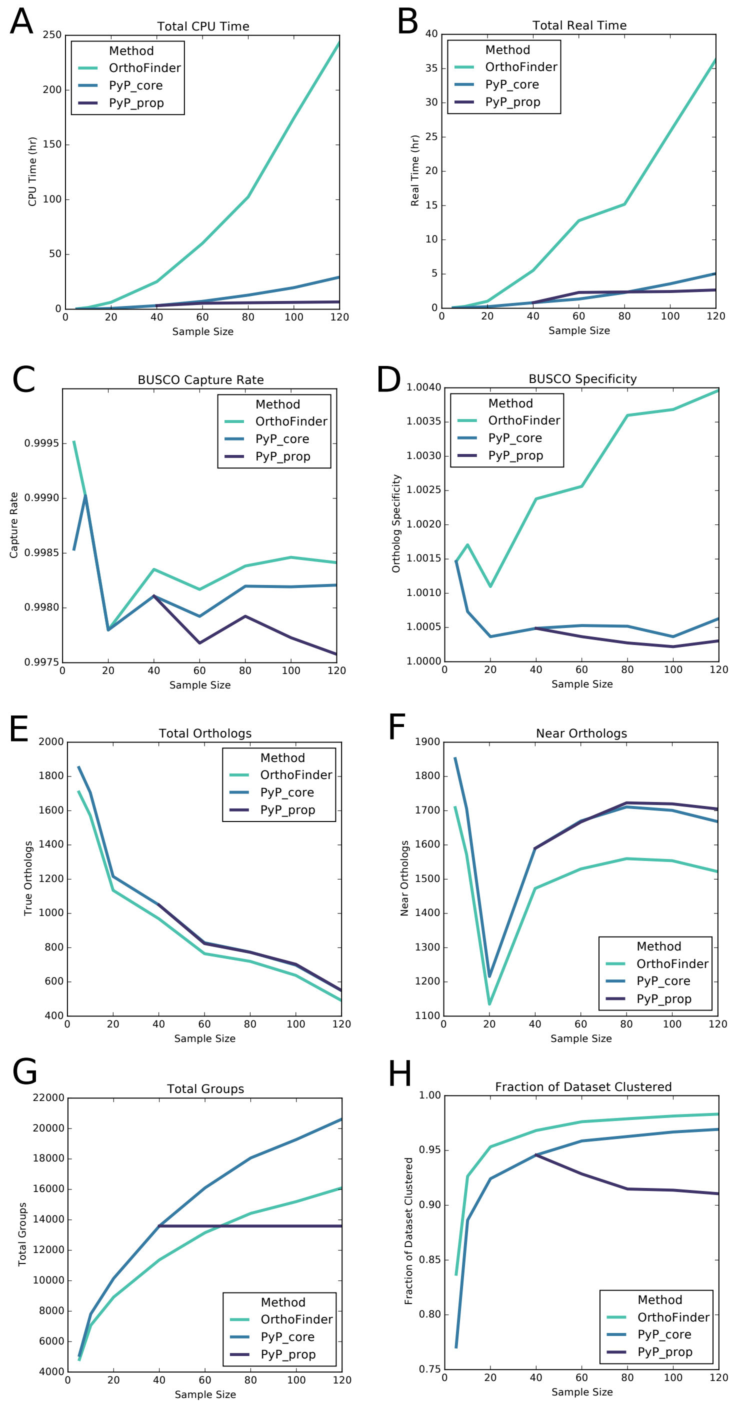


***Figure S3.*** Benchmarking both the core and propagation PyParanoid methodologies against OrthoFinder2. (**A**) Total CPU time and (**B**) total real time required for the three methodologies to be run on sample datasets. (**C**) Capture rate and (**D**) specificity of single-copy genes using the BUSCO database. (**E**) Total 'true' orthologs (exactly one per strain), (**F**) near orthologs (exactly one in >95% of strains), (**G**) total groups, and (**H**) fraction of the input dataset classified into groups (i.e. coverage).

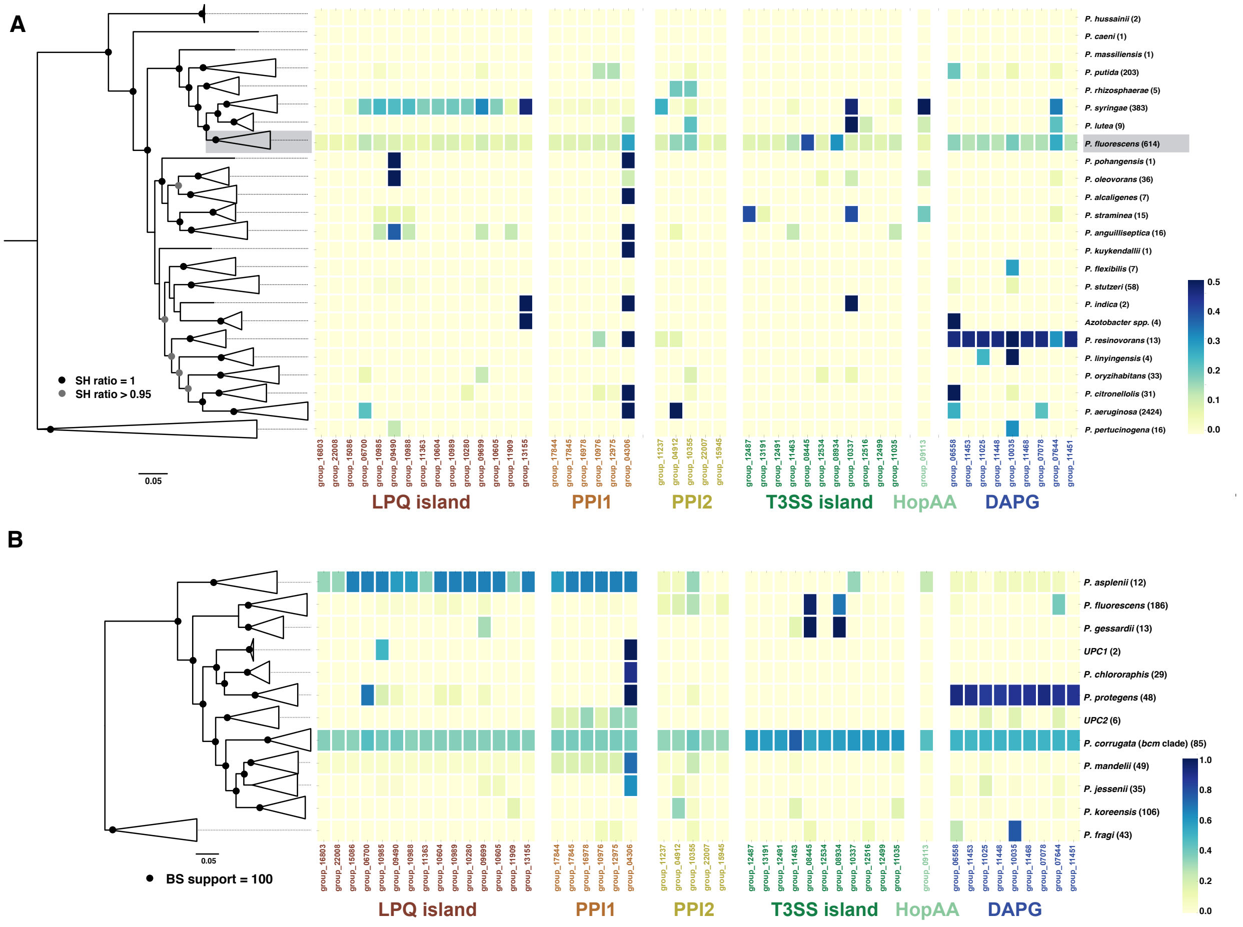


***Figure S4.*** Distribution of the 6 lifestyle-associated loci from across *Pseudomonas*. (**A**) shows the distribution of the loci within the *Pseudomonas* genus. Highlighted with a gray background is the *P. fluorescens* clade. (**B**) shows the distribution of the loci within the *P. fluorescens* clade. Taxa names and definitions have been previously described[25]. The names of all 6 loci are color-coded to be consistent with Figures 3 and 4 in the main text.

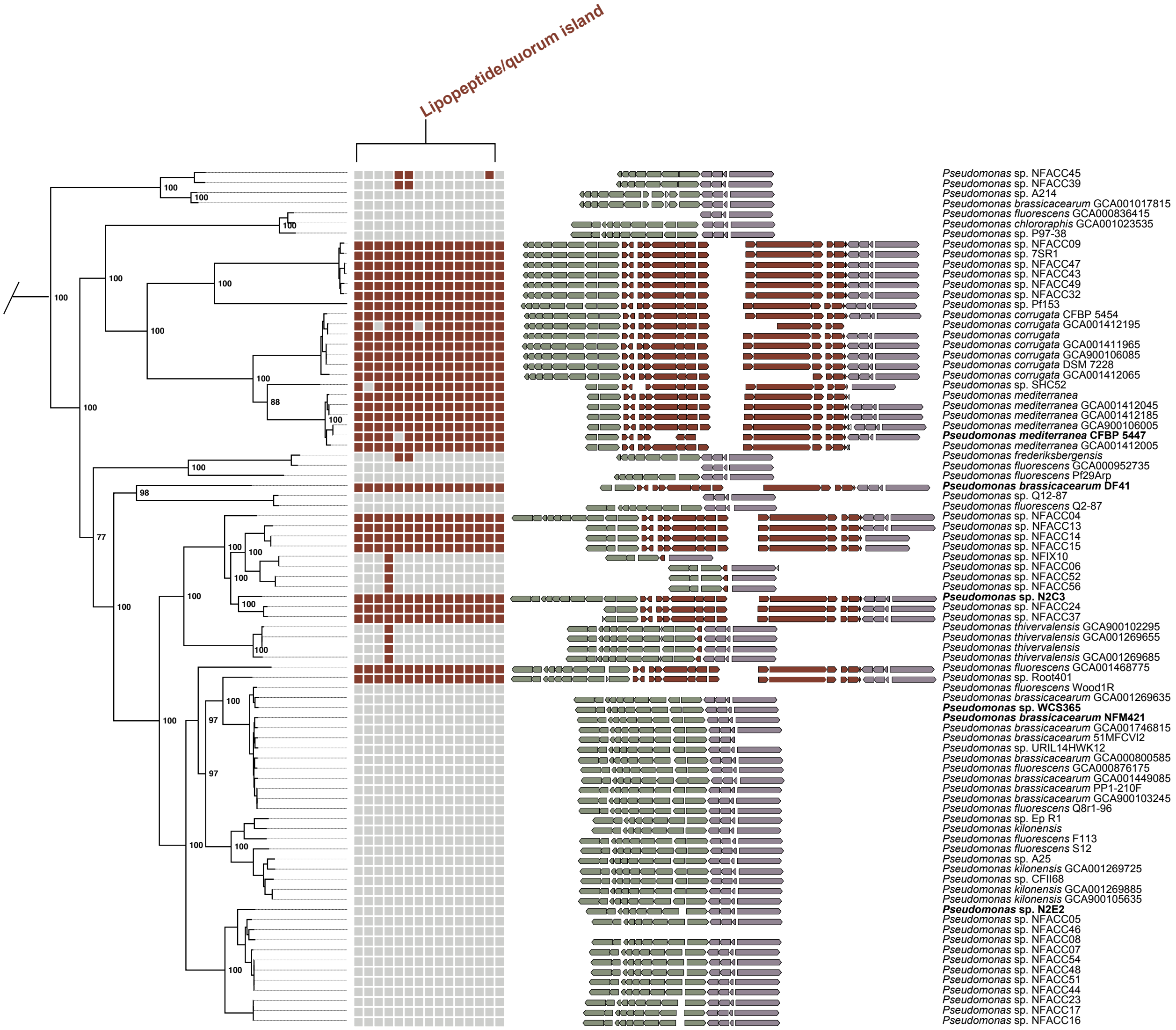


***Figure S5.*** Synteny plot showing the conservation of flanking regions for the LPQ island. The tree on the left side for all synteny plots is extracted from the *bcm* phylogeny from Figure 3 with the outgroup removed. Additionally, the presence (colored square) or absence (light gray square) for each PyParanoid homolog group associated with a given loci is depicted immediately to the right of the tree tips. The synteny plots are color coded to reflect conservation of a given integration site and ordered such that the loci of interest are oriented in the same direction. The green genes are exclusively found upstream of the LPQ island, whereas the purple genes are located downstream. Taxa with missing synteny plots do not have robust evidence of adjacent upstream and downstream regions, due to assemblies being too highly fragmented in the region surrounding the insertion site. The basic structure of this plot is conserved for Figures S7-S11, which show the conservation of the insertion site for the five other lifestyle-associated loci. Due to the size and repetitive nature of the LPQ island, the synteny plots for this region show only the ends of the island and not the interior region. Genome assemblies frequently break in the non-ribosomal peptide synthetase region, thus a contiguous DNA sequence representing the entire LPQ island can only be found in the 4 completed genomes with the island (*P. corrugata, P. mediterranea*, *P. brassicacearum* DF41, and *Pseudomonas* sp. N2C3).

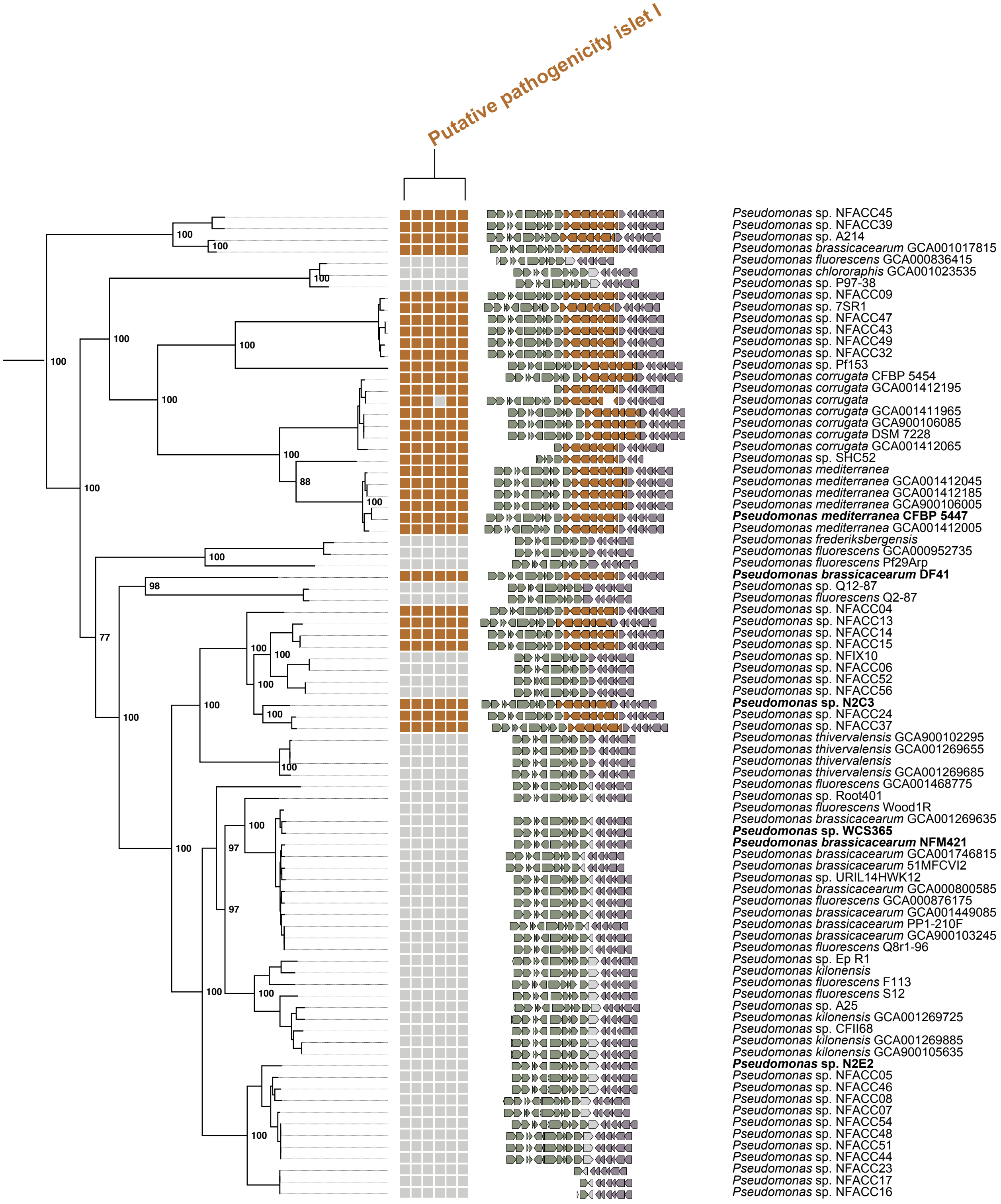


***Figure S6.*** Synteny plot showing the conservation of flanking regions for the putative pathogenicity islet I (PPI1). Layout described in Figure S6 caption.

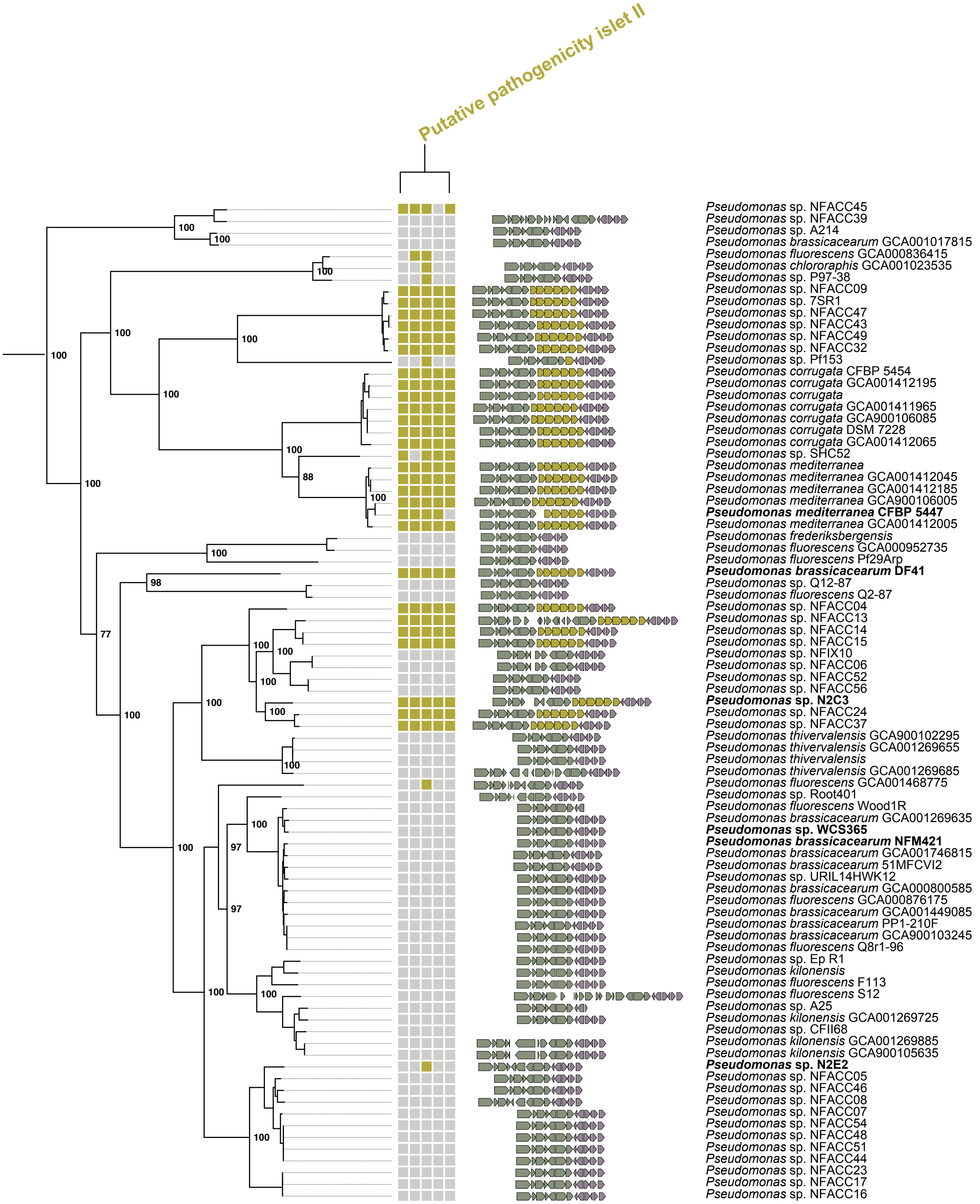


***Figure S7.*** Synteny plot showing the conservation of flanking regions for the putative pathogenicity islet II (PPI2). Layout described in Figure S6 caption.

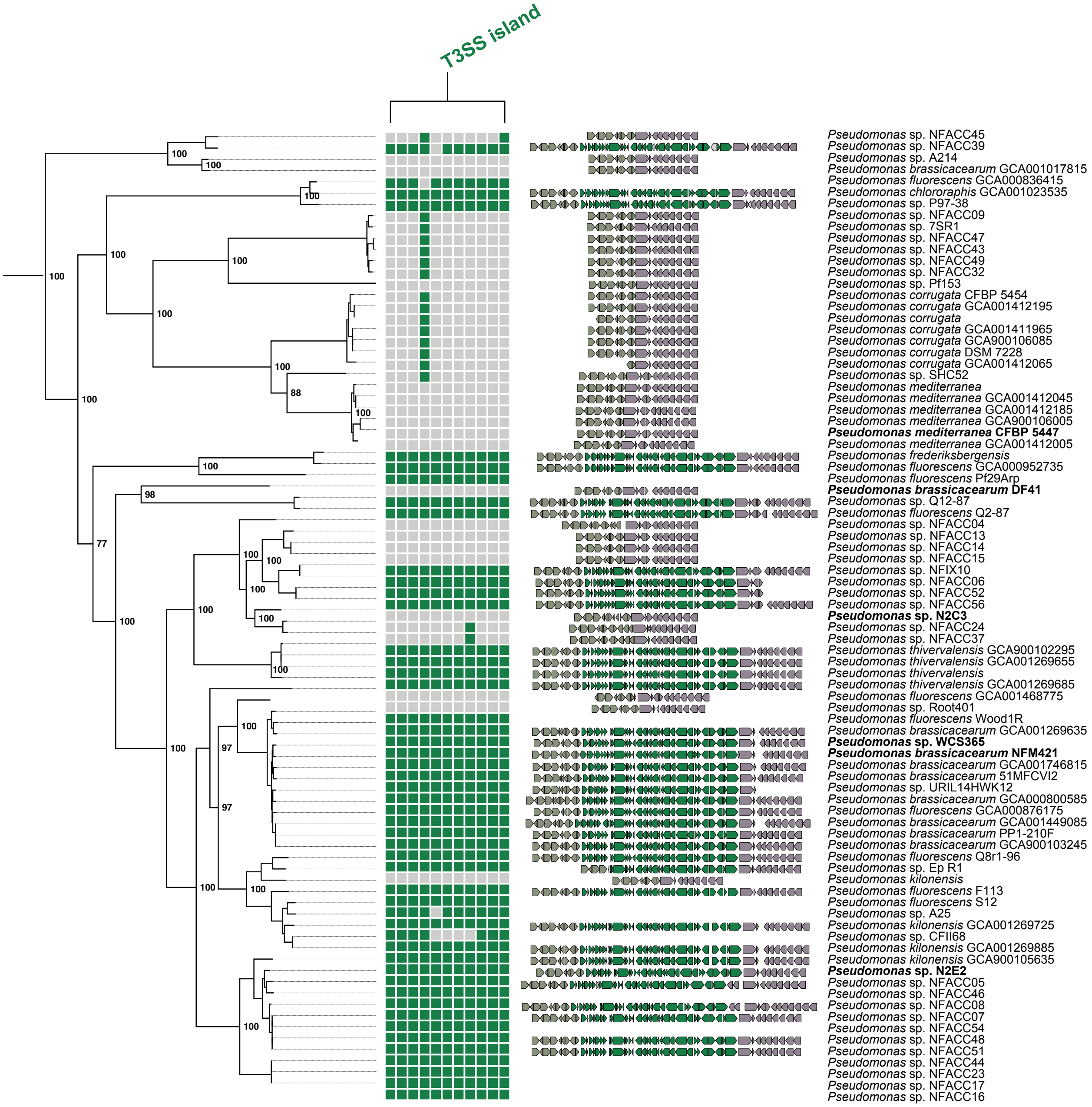


***Figure S8.*** Synteny plot showing the conservation of flanking regions for the type III secretion system (T3SS) island. Layout described in Figure S6 caption.

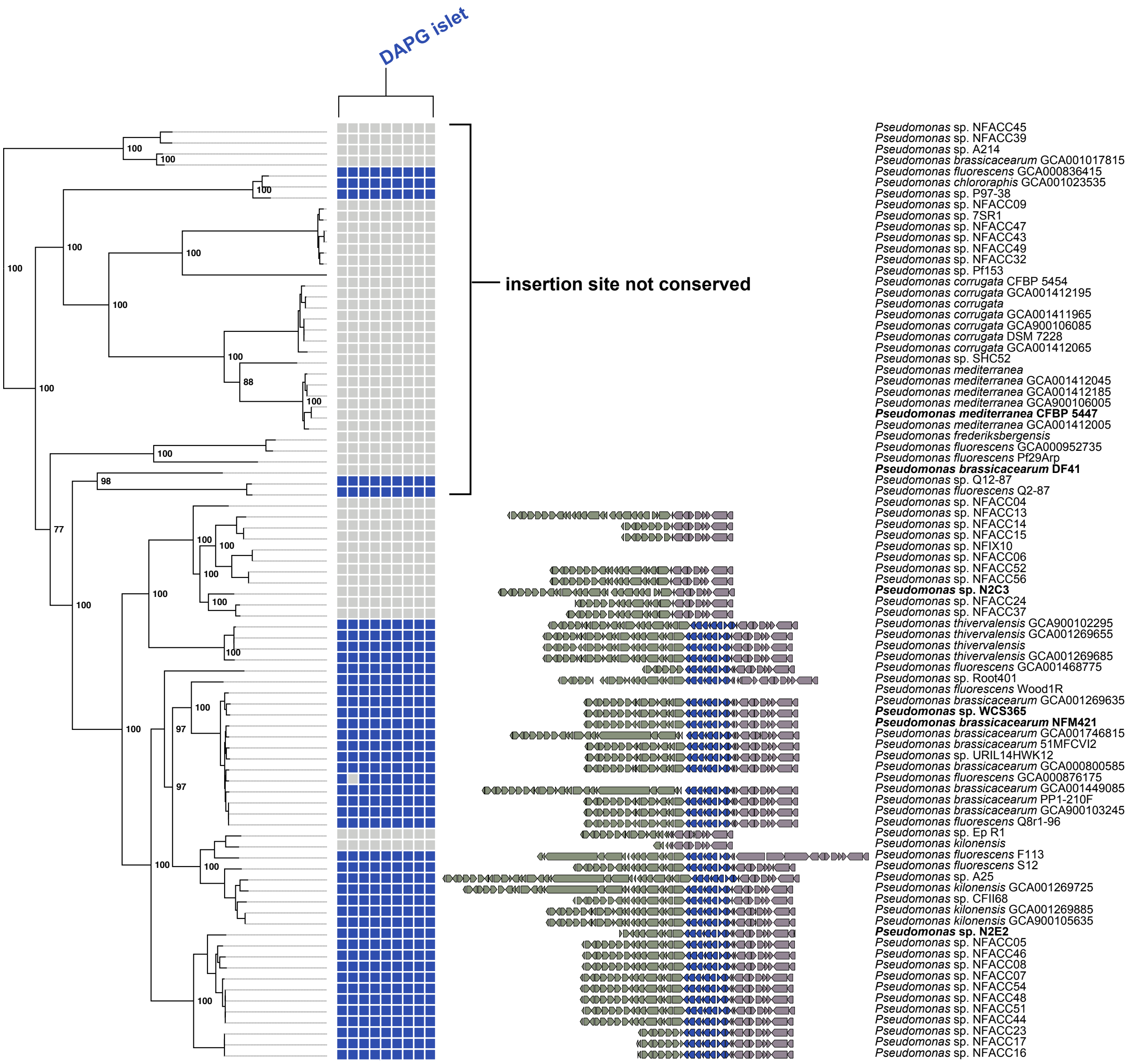


***Figure S9.*** Synteny plot showing the conservation of flanking regions for the diacetylphloroglucinol (DAPG) islet. Layout described in Figure S6 caption. Because the genomic region surrounding the putative DAPG insertion site was highly variable (see Figure 4B), we could only identify a conserved insertion site for the *P. brassicacearum* subclade since syntenic relationships in this region had highly diverged in the *P. corrugata*/*P. mediterranea* subclade.

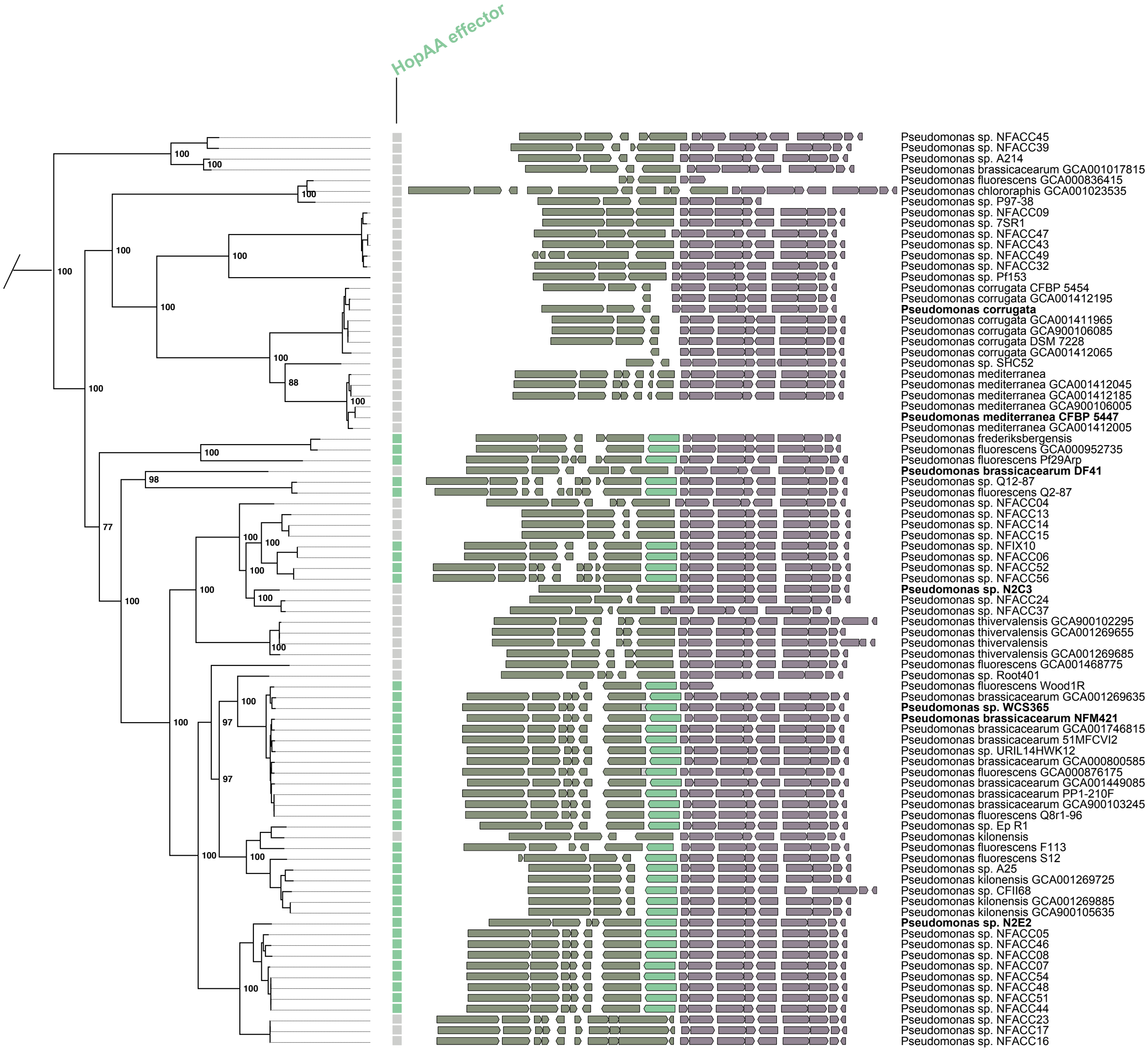


***Figure S10.*** Synteny plot showing the conservation of flanking regions for the type III secretion effector *hopAA* gene. Layout described in Figure S6 caption.

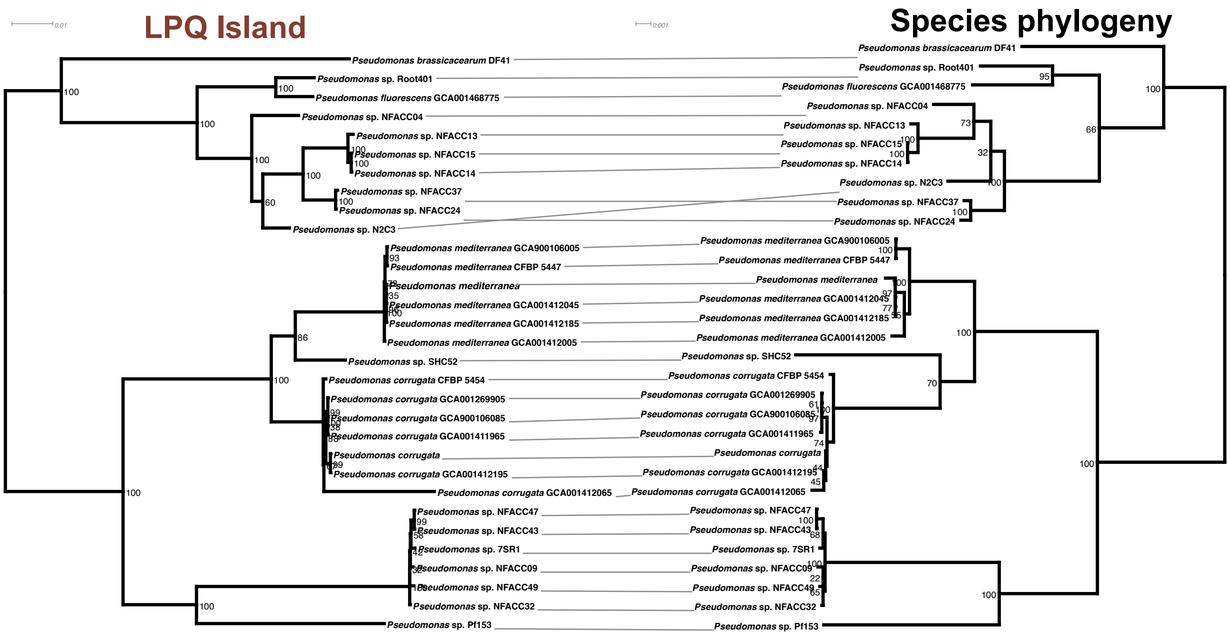


***Figures S11.*** Tanglegram showing the phylogeny of the LPQ island relative to the species phylogeny. The LPQ phylogeny was generated by aligning and concatenating 15 genes unique to the LPQ island. The species phylogeny was generated extracting only LPQ+ taxa from the alignment used to construct the *bcm* phylogeny in Figure 3. Note that there is only one incongruency between these trees and it is within a relatively closely-related clade, indicating that the evolution of the LPQ island was largely vertical and/or recombination was limited to closely related strains.

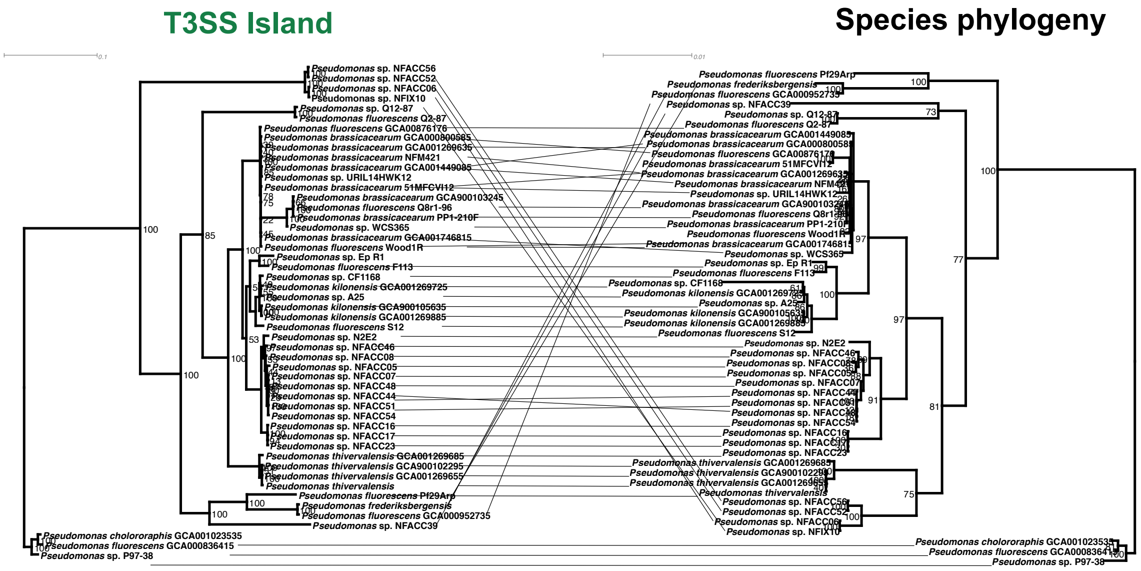


***Figure S12.*** Tanglegram showing the phylogeny of the T3SS island relative to the species phylogeny. The T3SS phylogeny was generated by aligning and concatenating 23 genes unique to the T3SS island. The species phylogeny was generated by extracting only T3SS+ taxa from the alignment used to construct the *bcm* phylogeny in Figure 3. Unlike the LPQ island, there are many examples of horizontal gene transfer and incongruencies between the T3SS phylogeny and the species phylogeny.

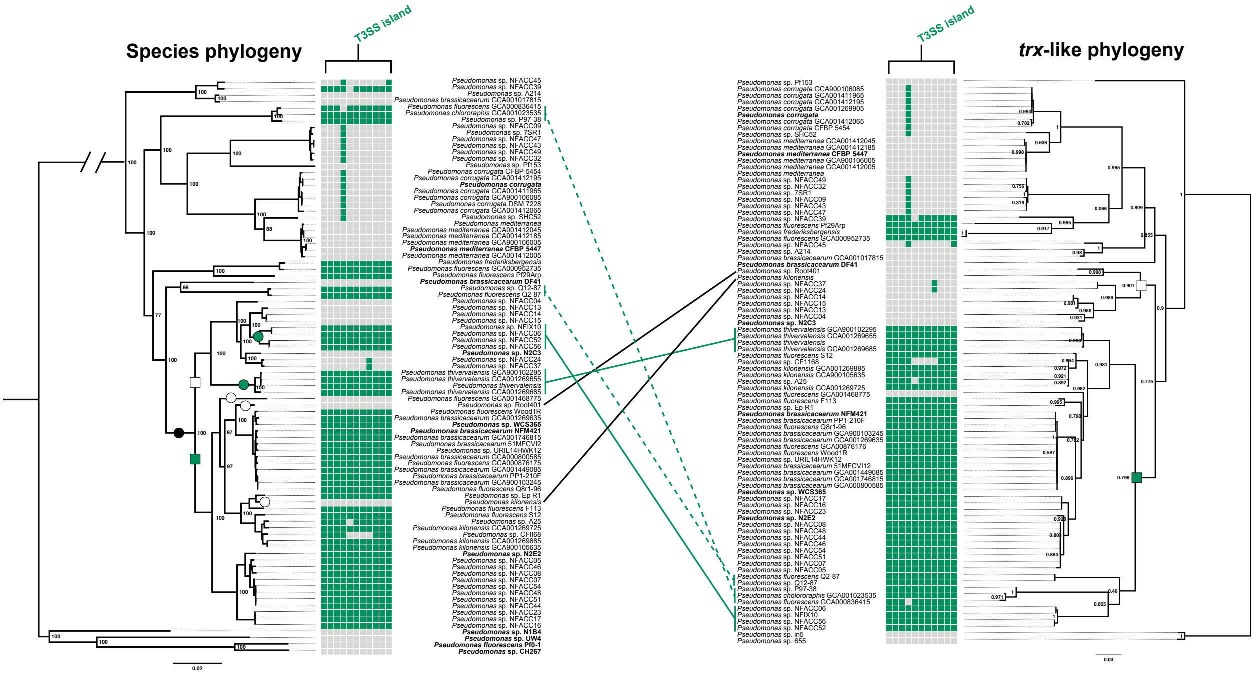


***Figure S13.*** Paired tree showing the phylogeny of the *trx*-like gene in relation to the *bcm* clade phylogeny. The *trx*-like gene phylogeny was based on a nucleotide alignment, while the *bcm* phylogeny from Figure 3 was used as the species phylogeny. Using parsimony to explain island presence and absence within the *P. brassicacearum* clade (black circle) based, we propose the clade diverged early into a commensal lineage with the T3SS island (green square) and a pathogenic lineage without the T3SS island (white square). Then, there were five putative recombination events leading to gain (green circles) or loss (white circles) of the T3SS island. Within the *trx*-like phylogeny, we identified two lineages that roughly corresponded to the putative lineages from the species tree (green and white squares). However, we identified six incongruencies between these pairs of predicted lineages. Four of these six incongruencies correspond to four of the five predicted island gain and loss events in our parsimonious model of T3SS island evolution. Two of these incongruencies represent island gain from a donor in the beneficial lineage (solid green lines) while two are island deletions from a donor in the pathogenic lineage (black lines); both of these events are diagrammed in Figure 4D. Moreover, there are two additional events associated with island gain outside of the *P. brassicacearum* clade (green dashed lines), indicating that homologous recombination of flanking regions is sufficient to explain most of the extant variation in the presence or absence of the T3SS island.

**Table Legends**

***Table S1.*** A list of strains used in this study as well as the isolation sites from the genomes used for *bcm* clade phylogenomics.

***Table S2.*** A list of primers used in this study.

***Table S3.*** Presence of the 15 unique lipopeptide island genes across *Pseudomonas* depicted in Figure 2F. While the lipopeptide biosynthesis genes are also found in many *P. syringae* strains, the entire complement of 15 genes is only found in strains from the *P. fluorescens* clade.

***Table S4.*** Association of loci in the *bcm* clade with the LPQ island. Correlation of homology groups with the LPQ island is based on the Pearson correlation across extant strains and is color coded (red indicates correlation with LPQ island presence, green with absence). The p-values for the 3 significance tests performed by treeWAS are also shown with p-values < 0.01 highlighted in green. The six loci chosen for further analysis are also noted and color coded based on the colors used in Figures 3, 4, and S5-S10.

***Table S5.*** Locus tags or accession numbers for the genes from the five islands and the *hopAA* gene in the *bcm* clade.
